# Neuropathy and neural plasticity in the subcutaneous white adipose depot

**DOI:** 10.1101/480095

**Authors:** Magdalena Blaszkiewicz, Jake W. Willows, Amanda L. Dubois, Stephen Waible, Cory P. Johnson, Kristen DiBello, Lila L. Lyons, William P. Breeding, Karissa B. Tilbury, Merilla Michael, Benjamin Harrison, Kristy L. Townsend

## Abstract

The difficulty in obtaining as well as maintaining weight loss, together with the loss of metabolic control in conditions like diabetes and cardiovascular disease, may represent pathological situations of inadequate neural communication between the brain and peripheral organs and tissues. Innervation of adipose tissues by peripheral nerves provides a means of communication between the master metabolic regulator in the brain (chiefly the hypothalamus), and energy-expending and energy-storing cells in the body (primarily adipocytes). Although chemical and surgical denervation studies have clearly demonstrated how crucial adipose tissue neural innervation is for maintaining proper metabolic health, we have uncovered that adipose tissue becomes neuropathic in various conditions of metabolic dysregulation. Here, utilizing both human and mouse adipose tissues, we present evidence of adipose tissue neuropathy, or loss of innervation, under pathophysiological conditions such as obesity, diabetes, and aging, all of which are concomitant with insult to the adipose organ and metabolic dysfunction. Neuropathy is indicated by loss of nerve fiber protein expression, reduction in synaptic markers, and less neurotrophic factor expression in adipose tissue. Aging-related adipose neuropathy particularly results in loss of innervation around the tissue vasculature. These findings underscore that peripheral neuropathy is not restricted to classic tissues like the skin of distal extremities, and that loss of innervation to adipose may trigger or exacerbate metabolic diseases. In addition, we have demonstrated stimulation of adipose tissue neural plasticity with exercise, cold exposure or neurotrophic factor treatment, which may ameliorate adipose neuropathy and be a potential therapeutic option to re-innervate adipose and restore metabolic health.

---

Since body weight regulation involves a precise balance between energy intake and energy expenditure, and requires coordination and control stemming from the central nervous system (CNS), the brain needs to adequately communicate with peripheral organs and tissues via peripheral nerves in order to maintain metabolic health. The CNS is able to control energy expenditure by signaling motivation to exercise or seek certain foods, driving sympathetic nervous system (SNS) activation of energy expending processes in the peripheral tissues of the body, and through receiving feedback from tissue sensory nerves, like those from adipose^1^. Sensory nerves in white adipose tissue (WAT) are thought to communicate the status of energy stores to the brain, in order to help regulate energy intake versus expenditure. In WAT, SNS activation serves to increase lipolysis, *de novo* adipogenesis, and a process termed ‘browning’, whereby uncoupling protein 1 (UCP1)-positive brown adipocytes appear in WAT depots and contribute to energy expenditure via thermogenesis^2^. In brown adipose tissue (BAT), SNS activation also drives non-shivering thermogenesis via activation of mitochondrial UCP1, which is a brown adipocyte-specific gene and required for this energy expending process in both classical and inducible/recruitable brown adipocytes^3,4^. During thermogenesis, BAT must utilize fatty acid fuels, which are both stored in the multilocular lipid droplets of brown adipocytes and are also obtained via circulating lipids that are released from WAT through lipolysis^5,6^.

Surgical and chemical denervation studies have demonstrated the importance of adipose tissue nerves, and denervation leads to a loss of proper metabolic control^3,7,8^. Denervation of WAT also leads to an increase in tissue mass and adipocyte cell number^9-11^. Therefore, the regulation of lipid stores in both BAT and WAT and the activation of energy expenditure through CNS-SNS communication are essential for proper body weight maintenance and metabolic health. It is not yet understood what other aspects of adipose tissue function are under neural control, and whether or not the nerves that innervate adipose tissue can modify their connections under physiological or pathophysiological conditions. Therefore, these questions regarding adipose nerve remodeling warrant further investigation.

The nerves that innervate white adipose tissues include numerous peripheral nerve subtypes, such as sensory, parasympathetic and sympathetic nerves ^12-14^. Although the number of studies assessing adipose innervation are increasing, further demonstrating the importance of brain-adipose communication^15-17^, it is still unclear which neurotransmitters and neuropeptides, aside from the well-studied norepinephrine, are synaptically released in adipose tissue and onto which receptor-expressing cell types. A better understanding of how the peripheral nerves in adipose tissue are regulated is important for the field, including differences in nerve plasticity between innervation of BAT and WAT, as well as sex differences in innervation density.

In other tissues, and in many organisms, peripheral nerves are appreciated as plastic (ie: able to undergo remodeling of neurites and synapses in response to stimuli), or neuropathic (dying-back in pathophysiological conditions). For example, with distal peripheral neuropathy nerves in the skin of distal extremities can die back through an unclear process, resulting in pain, loss of sensation, and severe discomfort. The process begins in the skin and moves inward. Neuropathy can be caused by aging, certain drugs (such as antibiotics and chemotherapy agents), or diabetes ^18-20^. Diabetic neuropathy is especially prominent, affecting over 50% of diabetic individuals, and often leads to limb amputation ^21,22^. Aging is associated with a loss of metabolic regulation and an increased propensity for diabetes^23-25^, and is independently associated with peripheral neuropathy^26^. The debilitating aspects of peripheral neuropathy are largely due to the inability to prevent or treat these conditions, and the inability to halt and reverse the neurodegeneration. Standard clinical approaches include pain management or glucose regulation (with diabetic neuropathy), but no therapies are currently approved to mitigate nerve death or to stimulate peripheral nerve re-growth or re-myelination. It is important to understand whether or not neuropathy can extend below the skin into underlying adipose tissue, which may exacerbate metabolic disease.

Given the close association between situations of metabolic dysregulation (aging, obesity, diabetes) and peripheral neuropathy, and the clear importance of adipose tissue innervation for metabolic homeostasis, we sought to determine if adipose tissue nerves also undergo neuropathy with these conditions. To do this, we assessed human adipose tissue samples across a range of ages and body mass indices (BMIs), and also utilized mouse models of aging and obesity/diabetes. We hypothesized that age and obesity/diabetes would be positively correlated with a loss of proper innervation of adipose tissues. We also sought to determine which interventions could stimulate re-innervation of adipose tissue in situations of adipose neuropathy, through a beneficial and physiological process of nerve plasticity.

## Methods

### Mice, Metabolic Phenotyping, and in vivo Analyses

#### Young/Aged Sedentary/Exercised cohort

Age-matched C57BL/6J male mice (Jackson Laboratory, Bar Harbor, ME; stock number *000664 Black 6*) were aged to 16 months. Body weight and adiposity were compared to age-matched C57BL/6J male mice at 10-12 weeks old.

#### Free running-wheel exercise

Young and aged animals were single-housed in running wheel cages that allowed free access to running, for a period of 7 days. Control (sedentary) animals were either single-caged with locked running wheels, or caged in pairs without a running wheel.

#### BTBR ob/ob

Male BTBR +/+ (WT) and *ob*/*ob* mutant (MUT) mice (Jackson Laboratory, Bar Harbor, ME; BTBR.Cg-*Lep*^*ob*^/WiscJ, stock number 004824) were fed a standard chow diet and aged to a minimum of 12 weeks of age, when they exhibit a robust phenotype including obesity, diabetes, and hyperglycemia ^27^. These animals (aged 12-28 weeks) were then euthanized after Von Frey Analysis of tactile allodynia (see details in Supplement).

### Human adipose tissue analyses

Human adipose samples (subcutaneous and omental) were obtained from the Boston Nutrition Obesity Research Center (BNORC) adipose tissue core. Biopsies of scWAT and omental adipose were taken from patients during elective surgery. All patients were either non-diabetic or pre-diabetic, with the exception of four diabetic individuals, 2 of whom were in the BMI cohort and 2 in the aged cohort (indicated by asterisks in data plots; see Suppl. Tables S1-3 for relevant patient data). Frozen and fixed tissue (from only a subset of patients) samples were obtained for analyses, including histology and western blotting.

### Whole depot imaging and analysis

Mouse inguinal subcutaneous adipose depots were removed intact and immunostained as described in Supplemental Materials. Entire depots were imaged for Fig. 4 with a 10x objective on a Leica TCS SP8 or DMI6000 confocal microscope (Leica Microsystems, Wetzlar, Germany), by tiling z-stacks of the entire depth of tissue. Images ranged between 542-3464 tiles per depot, the average lied at approximately 1000 tiles per depot. Tiles were individually Z-projected and background subtracted (using the rolling ball method). Processed tiles were then thresholded into binary images and skeletonized. To analyze arborization of adipose nerves, skeletons were assessed for the following parameters: innervation density, total number of branches, total skeleton length and tortuosity. Branches less than 4μm in length were excluded from the analysis. All analyses were performed in FIJI image analysis software. Arborization parameters were normalized to tissue weight. Additional analyses of whole depots are described in Supplement.

### Statistical Analysis

For all animal experiments, mice were randomized to treatment groups to ensure no difference in starting body weight. All plots represent mean +/-SEM. Statistical calculations were carried out in Excel or GraphPad Prism software (La Jolla, CA, USA), utilizing ANOVA, Linear Regression, or Student’s T-test as indications of significance (specified in Figure legends). Gene and protein expression data were normalized to a housekeeper and analyzed by either ANOVA or by Student’s t-test, two-tailed, using Welch’s correction when variance was unequal. Error bars are SEMs. For all figures, *p < 0.05, **p < 0.01, ***p < 0.001, ****p < 0.0001.

### Ethical Statement

All procedures and handling of animals were performed in accordance with the University of Maine’s Institutional Animal Care and Use Committee (IACUC), to comply with the guidelines of the PHS Policy on Humane Care and Use of Laboratory Animals, and Guide for the Care and Use of Laboratory Animals. Human tissue samples were obtained from the Boston Nutrition and Obesity Research Center (BNORC) and were de-identified prior to being provided to our laboratory. Tissues were collected by BNORC under their IRB-approved protocol.

### Data Availability

The datasets generated and/or analyzed during the current study are available from the corresponding author upon reasonable request. Any significant data generated or analyzed during this study thus far are included in this published article (and its supplementary information files).

*Additional Methods can be found in Supplemental Material.*

## Results

### Obese and diabetic mice display subcutaneous white adipose tissue (scWAT) neuropathy

In order to investigate whether obesity/diabetes leads to adipose tissue neuropathy, we used BTBR mice with the *ob/ob* leptin-deficient mutation (MUT). These animals develop severe obesity, type 2 diabetes, hypercholersterolemia, and insulin resistance, with significantly increased body weight and blood glucose by 8 weeks of age ^27^. Although disease progression is slower in females when compared to males^28^, both sexes eventually develop diabetes and peripheral neuropathy, with nerve conductance deficits and intraepidermal nerve fiber loss evident by 9 and 13 weeks, respectively ^27^. While this diabetic neuropathy has been well described in skin and paw, the effects on adipose innervation have not been assessed.

By 12 weeks of age, male BTBR MUT mice exhibited the expected increase in body weight when compared to BTBR +/+ wild type (WT) mice (Fig. 1a), which was concurrent with increased adiposity (Fig. 1b). In a pilot cohort that also included heterozygous (+/-HET) BTBR mice, only MUT animals displayed a trend for increased body weight and inguinal scWAT weight, for both males and females (Suppl. Fig. S1a). Thus, HET mice were not included in subsequent analyses. Consistent with previous reporting, neuropathy was detectable in BTBR MUT mice around 12 weeks of age via a Von Frey mechanical nociceptive assay (male mice, Fig. 1c; female mice, Suppl. Fig. 1b). This test determines tactile sensitivity in the skin of the hind paw, as an indirect measure of small fiber peripheral neuropathy. These data revealed that male mice displayed a stronger phenotype of peripheral neuropathy of extremities (Fig. 1c compared to Suppl. Fig. 1b), consistent with previous reports ^28^.

**Figure 1.**
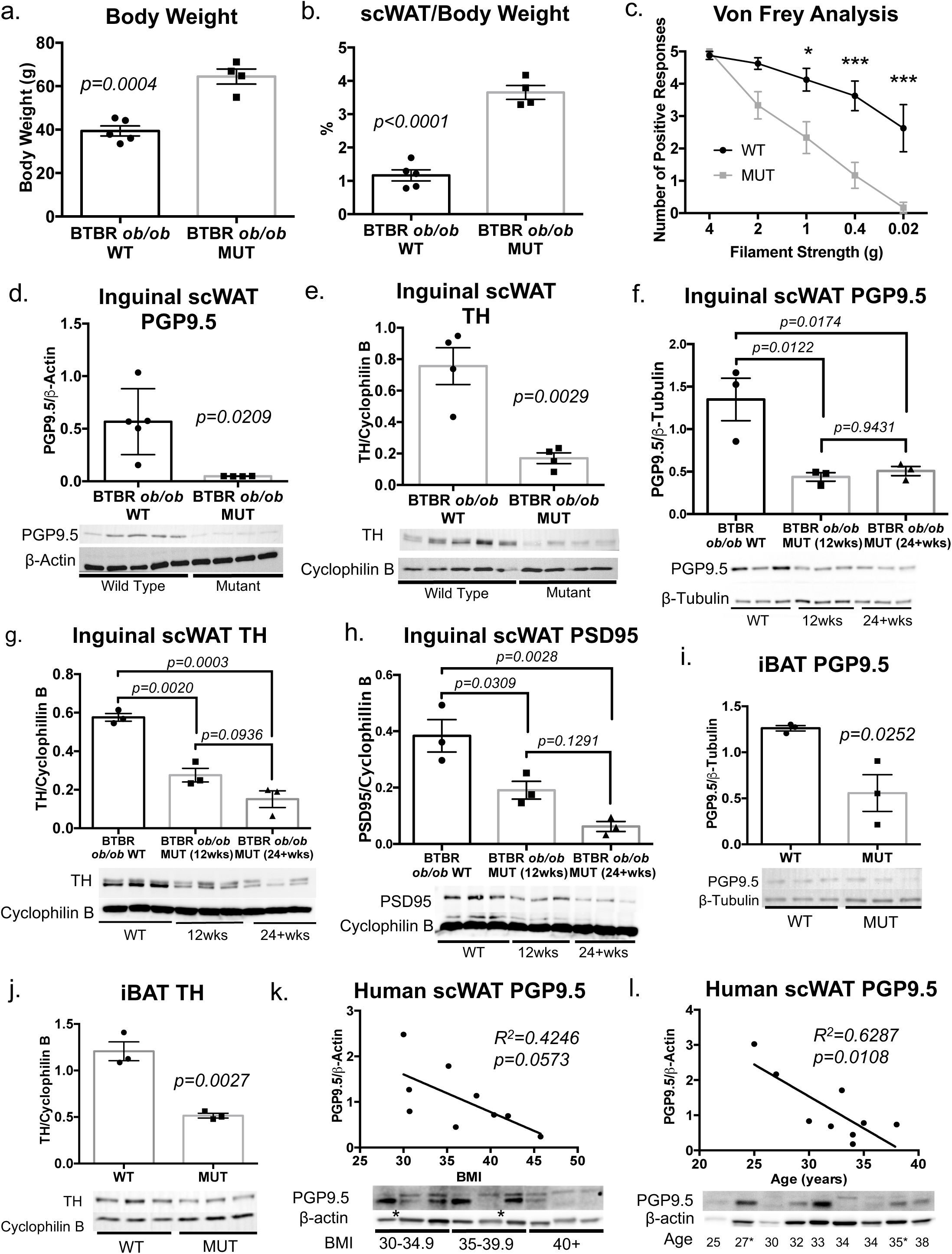
Obesity and diabetes led to white adipose tissue neuropathy. The *BTBR ob/ob* (MUT) model of obesity and diabetes was compared to *BTBR wild-type* (WT) for body weight measurements (a), and adiposity (b). Body and tissue weight data were analyzed by two-tailed Student’s T-Test. Von Frey tactile allodynia analysis was performed on MUT and WT animals to determine onset of peripheral neuropathy (c). Von Frey data was analyzed by ANOVA with Sidak’s *post hoc* test. For (a-c), all males; WT N=8, 12-20 weeks old; MUT N=6, 12-24 weeks old. Protein levels of PGP9.5 (d), as well as TH (e) in inguinal scWAT of the MUT and WT mice were measured by western blotting. For (e), lane 5 was excluded from analyses due to uneven resolution of housekeeper. For (d-e), all males; WT N=5, 12-20 weeks old and MUT N=4, 12-20 weeks old. Protein expression of PGP9.5 (f), TH (g) and PSD95 (h) in inguinal scWAT of 12-25 weeks old WT, 12 week old MUT and 24-28 week old MUT was measured by western blotting. All males, N=3 per group. Protein expression of PGP9.5 (i) and TH (j) in iBAT was measured in MUT mice and compared to WT littermate controls; for (i-j) all 12 week old males, N=3 WT/MUT. For all western blots, data were normalized to housekeeper proteins β-tubulin or cyclophilin B, band intensities were quantified in Image J, and analyzed using a two-tailed Student’s T-Test. Error bars are SEMs. In human scWAT, protein levels of pan-neuronal marker PGP9.5 were measured by western blotting; linear regression analysis was performed for assessment of normalized protein levels compared to body mass index (BMI) (k) or age (25-38 years old) (l). For (k-l) N=9 BMI cohort, N=9 Age cohort. For (k-l) * indicates individuals with diagnosed diabetes. The majority of human samples were females, see Suppl. Table S1-S3 for clinical details. Western blot data were normalized to β-actin, band intensity were quantified in Image J, and analyzed by two-tailed Student’s T-Test. Error bars are SEMs, *p < 0.05, **p < 0.01, ***p < 0.001, ****p < 0.0001.

The pan-neuronal marker protein gene product 9.5 (PGP9.5, also known as ubiquitin C-terminal hydrolase L1 (UCHL-1)), was also greatly reduced in inguinal scWAT of MUT mice (Fig. 1d; Suppl. Fig. S1c), indicating a loss of total nerve supply. BTBR MUT mice also displayed a marked reduction in sympathetic activation in inguinal scWAT as evidenced by a substantial decrease in tyrosine hydroxylase (TH) protein levels (Fig. 1e, Suppl. Fig. S1d). TH is the rate-limiting enzyme in catecholamine biosynthesis, including norepinephrine, the main sympathetic nerve neurotransmitter in WAT, and an indicator of SNS activation. After 12 weeks of age, the loss in total inguinal scWAT innervation remained consistent in MUT mice (Fig. 1f), but TH continued to decrease in MUT mice aged to 24 weeks (Fig. 1g). As loss of synaptic connections is another important factor in peripheral neuropathy, we measured protein expression of the synaptic protein postsynaptic density 95 (PSD95), and observed a reduction in PSD95 in 12-week-old MUT mice when compared to WT (Fig. 1h). PSD95 protein levels were also further decreased in 24-week-old MUT animals (Fig. 1h).

Because the energy expending interscapular brown adipose tissue (iBAT) is known to be highly innervated ^29,30^, we examined iBAT innervation in the BTBR mice. Protein expression of PGP9.5 revealed a significant decrease in total innervation of iBAT in BTBR MUT animals when compared to littermate controls (Fig. 1i). These finding were concurrent with a decrease in protein expression of TH in the iBAT of BTBR MUT mice indicating decreased sympathetic activation (Fig. 1j). iBAT gross morphology also revealed a marked increase in lipid accumulation, also called “whitening”, in both male and female mutant BTBR mice, but most prominently in male mice that exhibited the stronger neuropathy phenotype (Suppl. Fig. S1f). For BTBR MUT mice, iBAT gene expression showed a significant decrease in *ucp1*, whose activation by the SNS is necessary for thermogenesis to occur, along with reductions in the brown adipocyte markers *cidea* and *dio2,* despite no decrease in synaptic markers (*synapsin I, synapsin II, synaptophysin, psd95*), or other indicators of nerve health such as the Schwann cell marker *sox10* and neurotrophic factor brain derived neurotrophic factor (*bdnf*) (Suppl. Fig. S1g). The gene expression pattern in inguinal scWAT of BTBR MUT was opposite of what we observed in iBAT and included a coordinated trend for reduced expression of neural markers in MUT animals (Suppl. Fig. S1h). In a visceral WAT depot, peri-renal (prWAT), the gene expression pattern was similar to that of iBAT (Suppl. Fig. S1i). Together, these data indicate depot-specific differences related to innervation, and suggest that different mechanisms may be involved in maintaining nerve health in different adipose depots.

To determine whether peripheral neuropathy extended beyond adipose in BTBR MUT mice, we examined the neuromuscular junctions (NMJ). Immunofluorescent staining of NMJs revealed that BTBR MUT mice had fewer fully occupied NMJs in both the medial gastrocnemius (MG) and soleus (SOL) muscles when compared to WT littermates, accompanied by an increase in partially occupied junctions (Suppl. Fig. S2a-b). As there was no difference in unoccupied NMJs, neurodegeneration at the NMJ may be a slower neurodegenerative process with obesity and diabetes than in skin and underlying adipose.

We next attempted to visualize the changes to inguinal scWAT innervation of BTBR MUT animals within the whole depot. This proved technically difficult to accomplish using epifluorescent or standard confocal microscopy, due in part to the fibrotic nature of obese adipose tissue. However, with 2-photon microscopy it was possible to visualize structural changes within scWAT of BTBR MUT animals (Suppl. Fig. S2c). Immunofluorescent staining for PGP9.5 was performed on whole inguinal scWAT tissue, 2-photon fluorescence imaging was used to image the immunofluorescent staining of PGP9.5 (green) and Second Harmonic Generation (SHG) imaging of collagen fibers with a label-free method (Suppl. Fig. S2c). While collagen is abundant in both WT and MUT BTBR animals, a greater degree of colocalization between collagen fibers and nerves is visible in BTBR MUT animals (Suppl. Fig. S2c, yellow color), likely contributing to our inability to image the adipose nerves successfully in this mouse model.

### White adipose tissue from obese humans exhibits neuropathy

Based on our findings in the BTBR mouse model we sought to determine if neuropathy exists in adipose tissue of obese humans. We investigated degree of innervation in human scWAT (and omental WAT) obtained from individuals who underwent elective surgery. Protein expression of the pan-neuronal marker PGP9.5 revealed a decreasing trend with increasing BMI in human scWAT adipose tissue (Fig. 1k), despite no difference in adipocyte diameter (Suppl. Fig. S3a, left panel), indicating that adipose neuropathy also exists in human tissues. Since we were unable to obtain an entire WAT depot from humans, in order to determine if innervation status was confounded by cell size, adipocyte diameter was quantified for the human samples and revealed no differences that may confound these analyses. In addition, for all innervation analyses, western blots are always controlled by equal protein loading for each sample and normalization to a housekeeping protein. For mouse, entire inguinal scWAT depots were homogenized to analyze the total level of innervation, however, this was not possible for human samples as we could only obtain a small biopsy. Therefore, normalized protein expression was plotted against average cell size for each corresponding sample, which showed no correlation (Suppl. Fig. S3b). Interestingly, post-translational modification of PGP9.5 appears to be restricted to scWAT of humans (Fig. 1k-l) since we did not observe multiple band sizes in any of our mouse models.

We also assessed degree of innervation in human omental adipose tissue, and found no significant difference in innervation with increasing BMI (Suppl. Fig. S3c), suggesting that adipose neuropathy may be restricted to scWAT depots. This fits with our data from mouse as well. However, due to the limited number of omental adipose samples available, along with the narrow range in BMI, further assessments may be warranted to confirm this observation. Nevertheless, these data show that human adipose tissue undergoes neuropathy with obesity similar to the mouse model.

### Aging leads to white adipose tissue neuropathy in humans and mice

Since adiposity increases with age concordant to increased risk of developing type 2 diabetes, we sought to determine if there is a link between adipose neuropathy and aging. When investigating the relationship between adipose innervation and age in humans, linear regression analysis revealed a significant decrease in protein levels of the pan-neuronal marker PGP9.5 with increasing age in inguinal scWAT (Fig. 1l) despite no difference in adipocyte size with no correlation between average cell size and protein expression (Suppl. Fig. S3a-b, right panels). Again this finding did not extend to omental WAT (Suppl. Fig. S3d).

To further explore this relationship, we employed an aged mouse model. Male C57BL/6J mice were aged to 16 months, and the state of their adipose innervation was compared to young mice at 10-12 weeks old. Protein expression of PGP9.5 in inguinal scWAT was decreased in 16 month old aged mice when compared to young mice at 10-12 weeks of age (Fig. 2a), together with a trend for decreased TH levels (Fig. 2b). Consistent with the metabolic consequences of aging, total body mass of aged mice was significantly greater than that of young animals, however, there was no significant difference in quadriceps muscle weight or adipose depot weight, although inguinal scWAT and perigonadal (pg)WAT depot weight did display a trend to be higher at this age (Suppl. Fig. S4a).

**Figure 2.**
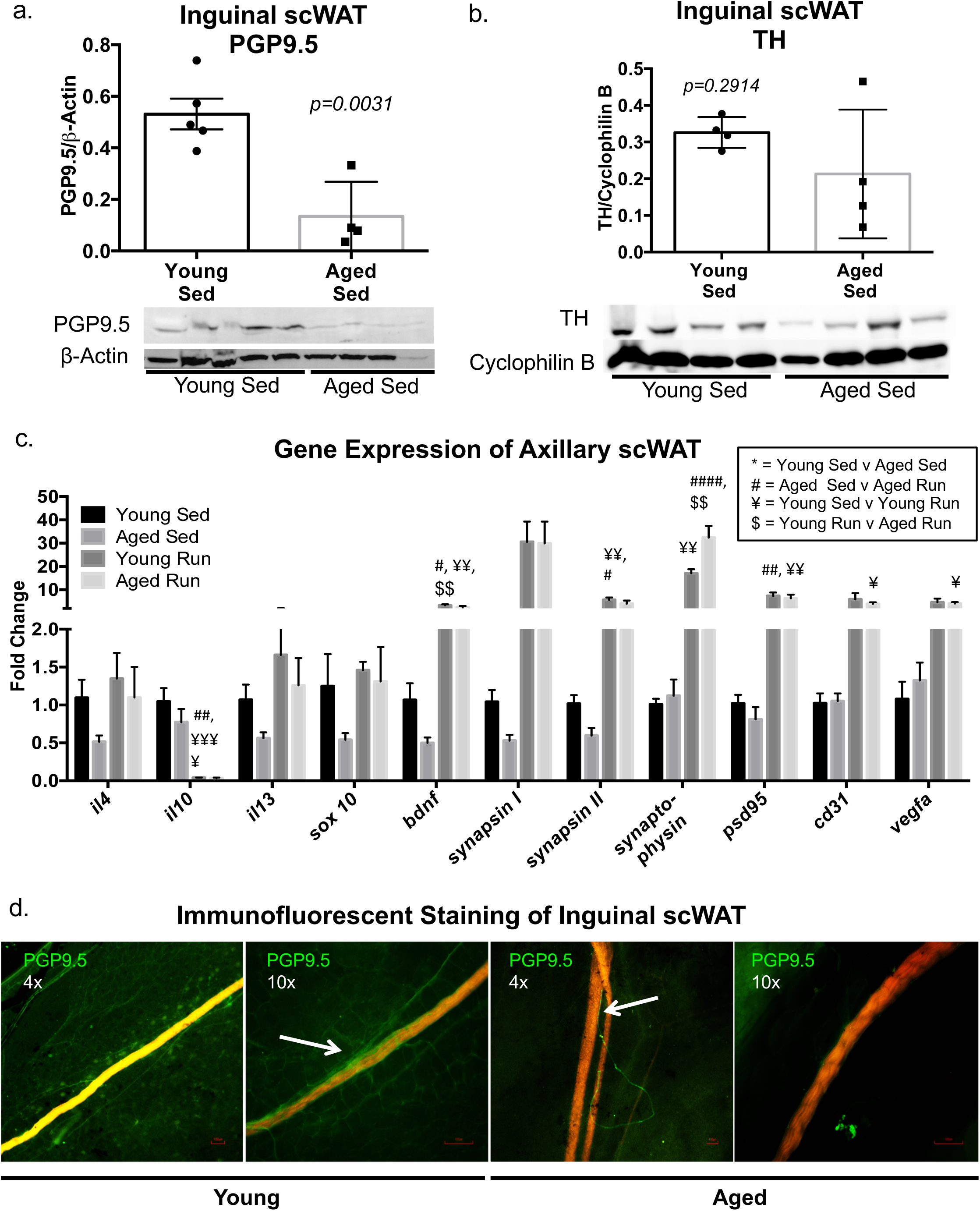
Aging is associated with adipose tissue neuropathy. In young (10-12 weeks old) and aged (16 months old) sedentary male C57BL/6J mice, western blotting was used to measure protein expression of PGP9.5 in inguinal scWAT for assessment of total innervation (a), while TH was used to measure sympathetic activation (b). Protein expression was normalized to either β-actin or cycophilin b, band intensity was quantified in Image J and analyzed by two-tailed Student’s T-Test; N=4 for young and aged groups. Gene expression of anti-inflammatory markers *il4* and *il13, il10*, neurotrophic factor *bdnf*, synaptic markers (*synapsin I and II, synaptophysin, psd95*), as well as vascular markers (*cd31, vegfa*) was measured by qPCR in axillary scWAT of young and aged mice (c). For young and aged sedentary animals N=4, for young and aged exercised (run) mice, N=5 per group. Gene expression data expressed as fold change normalized to the housekeeping gene *cyclophilin*, was analyzed by two-way ANOVA, with Uncorrected Fisher’s LSD post hoc test, and significance denoted with * for young sedentary versus aged sedentary; # for aged sedentary versus aged run; ¥ for young sedentary versus young run; and $ for young run versus aged run. Whole depot immunofluorescent imaging of inguinal scWAT from sedentary animals for total innervation (PGP9.5 in green) and vasculature (red/orange; autofluorescence) was performed, arrows indicate innervation around vasculature (d). Images captured at 4x and 10x, and are representative of N=4 mice analyzed per group. Error bars are SEMs, *p < 0.05, **p < 0.01, ***p < 0.001, ****p < 0.0001.

Since a state of chronic inflammation increases tissue dysfunction and is known to damage nerves ^31^, we measured levels of anti-inflammatory cytokines which are involved in inflammatory resolution ^32^ in young and aged sedentary mice. A non-significant reduction in the anti-inflammatory cytokines *interleukin 13* (*il13*), *interleukin 4* (*il4*), but not *interleukin 10* (*il10*), was observed in the 16 month old mice when compared to younger mice in the sedentary state (Fig. 2c), suggesting an altered inflammatory homeostasis in aged adipose tissue. Gene expression for presynaptic proteins (*synapsin I and II*), the neurotrophic factor *bdnf*, which supports growth and survival of nerves, as well as the Schwann cell marker *sox10*, showed a coordinated trend for reduction in aged mice as well (Fig. 2c).

Considering that nerves and vasculature are often linked, we measured gene expression of vascular markers (*cd31, vegfa*) in young and aged sedentary mice and saw no difference with respect to age (Fig. 2c). We took advantage of vascular autofluorescence^33^ to visualize adipose blood vessels in combination with immunofluorescent staining of nerves with a new whole-depot 3D microscopy technique developed in our lab (see Supplement). This imaging revealed that aged mice exhibit an overall decrease in innervation with a striking loss of innervation around vasculature within their inguinal scWAT (Fig. 2d). In young mice, autofluorescent blood vessels (red/orange) are clearly shrouded by fine PGP9.5-expressing nerves (green, Fig. 2d, left panels), but by 16 months of age, the adipose vasculature shows a striking loss of this neuronal sheathing (Fig. 2d, right panels). Taken together, these data reveal that loss of proper adipose innervation may precipitate aging-related metabolic dysfunction.

### Exercise attenuates age-related adipose neuropathy

Exercise has been shown to increase circulating levels of the nerve survival factor BDNF ^34^. We therefore exercised young and aged mice on running wheels for 7 days and assessed the status of their adipose innervation. Two-way ANOVA analysis of gene expression revealed an interaction of exercise and age only for *synaptophysin*, with exercise (p<0.0001) having a stronger effect than age (p=0.0213) (Fig. 2c). Post-hoc analyses revealed an exercise effect to increase *bdnf, synapsin II* and *psd95*, indicating an improvement in innervation. This occurred despite no changes in total body mass or adiposity before and after the short exercise intervention (Suppl. Fig. S4a). This increase in innervation was observed despite no difference in anti-inflammatory cytokines *il4, and il13* in both young and aged animals after exercise (Fig. 2c). Exercise also increased gene expression of vasculature markers *cd31* and *vegfa*, but only in young animals (Fig. 2c), indicating that physical activity can increase both vascular and nerve supply to adipose, but perhaps is blunted with advancing age.

Protein levels of both PGP9.5 and TH in inguinal scWAT of young exercised mice were significantly increased when compared to young sedentary mice (Fig. 3a-b), indicating that neurite outgrowth has likely taken place, as well as sympathetic nerve activation. Exercise also resulted in a trend for increased protein expression of PGP9.5 and TH in aged animals (Fig. 3c-d), although this did not reach significance, further supporting a blunted effect with age. Direct comparison of the effect of exercise on inguinal scWAT innervation of young versus aged animals revealed no significant difference in response based on age, as both young and aged mice showed relatively similar levels of PGP9.5 and TH expression (Suppl. Fig. S4b-c). The effects of exercise on adipose tissue did not extend to BAT. Protein expression of PGP9.5, TH, and PSD95 in BAT did not differ between young sedentary and exercised groups (Suppl. Fig. S4d-f), nor did gene expression of nerve and synaptic markers (Suppl. Fig. S4g).

**Figure 3.**
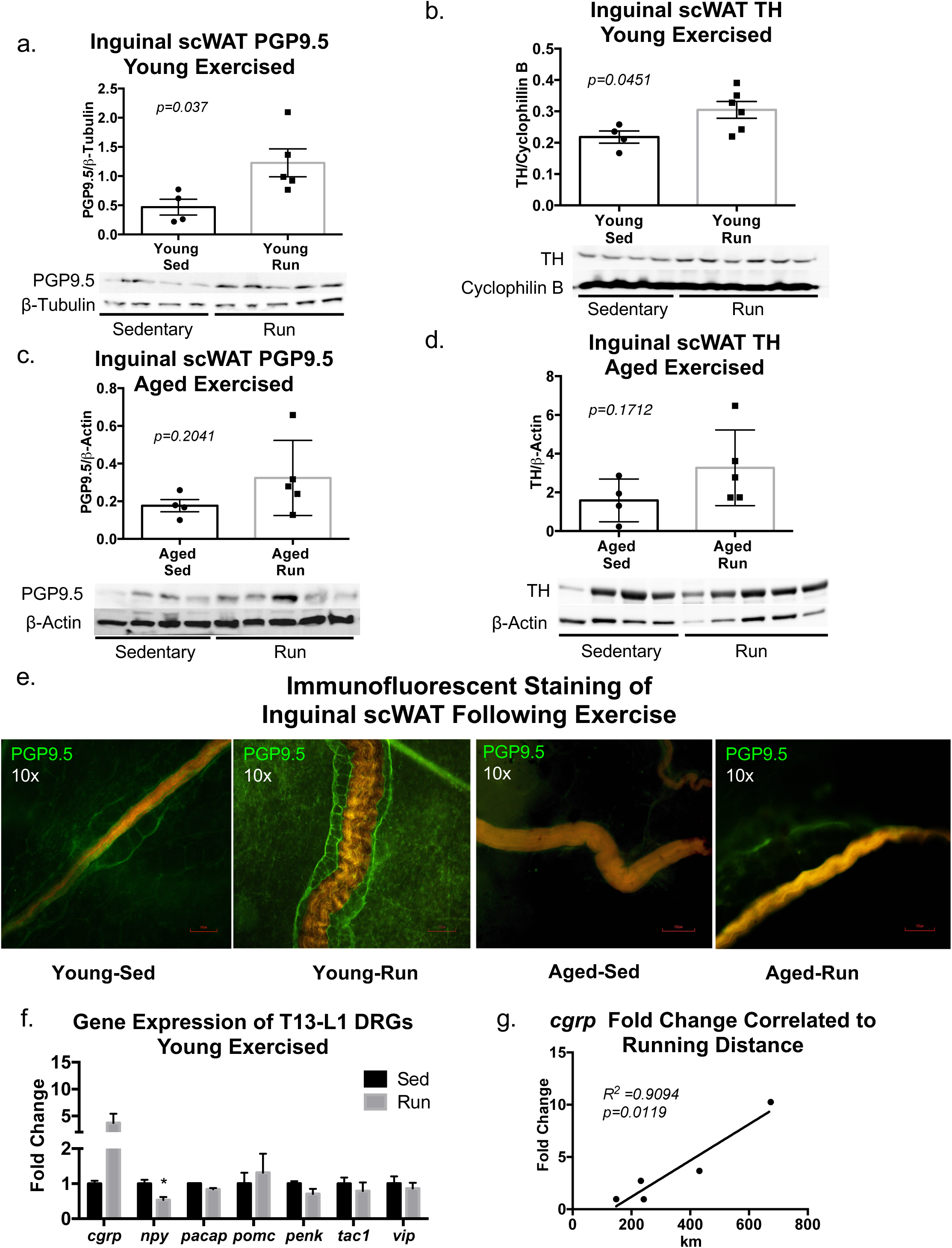
Exercise increases adipose innervation in young mice and attenuated loss of age-related adipose innervation. Young (10-12 weeks old) and aged (16 months old) male C57BL/6J mice were placed in running-wheel cages for 7 days with continuous access to a running wheel (run). To assess exercise effects on adipose innervation in young mice, protein expression in inguinal scWAT was measured by western blotting with PGP9.5 as a marker of total innervation (a), and TH as an indicator of sympathetic activation (b). Protein expression was normalized to either β-tubulin or cyclophilin b, band intensity was quantified in Image J and analyzed by two-tailed Student’s T-Test; N=4 for young sedentary and N=5-6 for exercised groups. Protein expression of PGP9.5 (c) and TH (d) in inguinal scWAT of aged sedentary and exercised mice was also determined by western blotting. Protein expression was normalized to β-Actin, band intensity was quantified in Image J and analyzed by two-tailed Student’s T-Test; N=4 for aged sedentary animals, N=5 for aged exercised (run) animals. Whole depot nerve and vasculature imaging of inguinal scWAT from exercise (run) animals was performed by combining immunostaining for PGP9.5 (green) with autofluorescence of vasculature (red/orange); images were captured at 10x (e). Images are representative of N=5 mice analyzed per group. Young (12-15 weeks old) male C57BL/6J mice were placed in running-wheel cages for 7 days with continuous access to a running wheel (run), control (sed) animals were placed in cages with a locked running wheel. Gene expression of neuropeptides in the T13-L1 DRGs was measured by qPCR, expressed as fold change normalized to the housekeeping gene *cyclophilin*, and analyzed by Student’s T-test (f). Linear regression analysis was performed for *cgrp* to assess the effect of running on changes in gene expression (g). Error bars are SEMs, *p< 0.05, **p < 0.01, ***p < 0.001, ****p < 0.0001.

Exercise did not appear to have a specific effect on the vasculature innervation, but the levels of PGP9.5 around vasculature had high intra-individual variability (Fig. 3e). In another approach to assess innervation of blood vessels, immunostaining of PGP9.5 was combined with isolectin staining of vasculature in inguinal scWAT. Innervation of blood vessels that measured 50 μm or more in diameter was evaluated (Suppl. Fig. S5a-b). We found no difference in either percentage of blood vessels innervated (Suppl. Fig. S5a, left panel) or number of nerves per blood vessel (Suppl. Fig. S5a, right panel), between sedentary and exercised groups.

In order to determine changes to sensory nerves innervating WAT, we assessed gene expression in the T13-L1 dorsal root ganglion (DRG), which is known to innervate inguinal scWAT^35,36^ (Fig. 3f). The following neuropeptides were measured with or without exercise intervention: calcitonin gene-related peptide (CGRP), neuropeptide Y (NPY), pituitary adenylate cyclase-activating polypeptide (PACAP), pro-opiomelanocortin (POMC), proenkephalin (PENK), substance p (Tac1), and vasoactive intestinal peptide (Vip). We found that mRNA expression of *npy* was decreased in exercised mice (Fig. 3f). The effect of exercise on *cgrp* expression was highly variable, but linear regression showed that this neuropeptide increased with relation to amount of running the animal performed (Fig. 3g), which fits with reports of increased *cgrp* expression in neurons of the T13/L1 ganglia upon inguinal scWAT cold induced ‘browning’ ^36^.

### Optimization of whole depot imaging of adipose innervation

In order to gain a comprehensive understanding of the neural network within inguinal scWAT we explored various immunostaining methods and microscopy techniques. Immunostaining of standard paraffin embedded 7uM tissue sections (Suppl. Fig. S6a) provides incomplete information, since nerve fibers are bisected and appear only as puncta. ScaleView-A2 tissue clearing method allowed for visualization of intact inguinal scWAT depots of a TH-reporter mouse (Suppl. Fig. S6b) and revealed region specific increases of sympathetic activation in 7 day cold exposed mice, when compared to animals that were cold exposed and then rewarmed for 4 weeks. Confocal microscopy with depth coding of sucrose-cleared inguinal scWAT revealed that nerves penetrate the inguinal scWAT depot and are not merely running along the surface (Suppl. Fig. S6c). Although tissue-clearing methods allowed for an intact depot without flattening and distortion of tissue boundaries, tissues (especially from 80g BTBR MUT mice) were too large for lightsheet microscopy, and too thick for standard confocal.

To overcome these limitations, a method was developed which allowed for whole depot visualization by flattening the tissue between two large slides, followed by acquisition of confocal tiled z-stack images that could be digitally reconstructed to present innervation of the entire depot, *<u>without the need for tissue clearing or lightsheet microscopy</u>*. Inguinal scWAT depots were flattened (70-100μm thickness), autofluorescence was blocked with Sudan Black staining, and entire depots were immunostained with PGP9.5. This technique revealed that individual adipocytes were surrounded by neurites, forming a honeycomb-like pattern throughout the majority of the tissue (Suppl. Fig. S6d); while larger myelinated nerve bundles could also be visualized (Suppl. Fig. S6d, right image). Synapses in adipose tissue also appear to occur directly in the stromal vascular fraction (SVF) (Suppl. Fig. S6e). In some tissues, heparin can be used to wash out erythrocytes from vasculature when its autofluorescence is undesired (Suppl. Fig. S6f).

We also compared the use of antibodies to nerve reporter mouse strains. Inguinal scWAT from *Na*_*v*_*1.8-Cre x tdTomato* reporter mice, which marks sensory nerves, was costained with PGP9.5, and results were compared to inguinal scWAT from C57BL/6 mice immunostained with Na_v_1.8 and PGP9.5 antibodies (Suppl. Fig. S7a). Non-specific staining with the Na_v_1.8 antibody was evident in the lack of co-localized signal with the pan-neuronal marker PGP9.5. Conversely, when we compared tissues from a sympathetic reporter mouse line (*TH-Cre x ROSA YFP*) to TH antibody, there was a much more robust signal from the reporter mouse tissue, indicating that the reporter may be picking up more nerve fibers than the antibody (Suppl. Fig. S7b).

### Adipose tissue undergoes remodeling with cold exposure

In addition to exercise as a means to promote adipose tissue neural plasticity, we found that cold stimulation also increased adipose tissue innervation, as protein levels of PGP9.5 went up in scWAT after 3 days of cold exposure compared to tissues from mice housed at room temperature or thermoneutrality (Fig. 4a). To examine the adipose nerve network under this innervation promoting condition, adult mice maintained at room temperature (RT) were compared to littermates that were either cold exposed for 10 days or cold exposed for 10 days and rewarmed for 1 week. Inguinal scWAT depots were immunostained with the pan-neuronal marker β-3-tubulin. At room temperature, a diffuse pattern of innervation can be discerned (Fig. 4b, left panel), with larger nerve bundles converging around the subiliac lymph node (white arrow), similar to the staining pattern we observe with the other pan-neuronal marker, PGP9.5. The posterior end (right of subiliac lymph node in image) of the tissue did not appear to have as much innervation as the anterior portion (left of subiliac lymph node in image) or as the area surrounding the subiliac lymph node.

**Figure 4.**
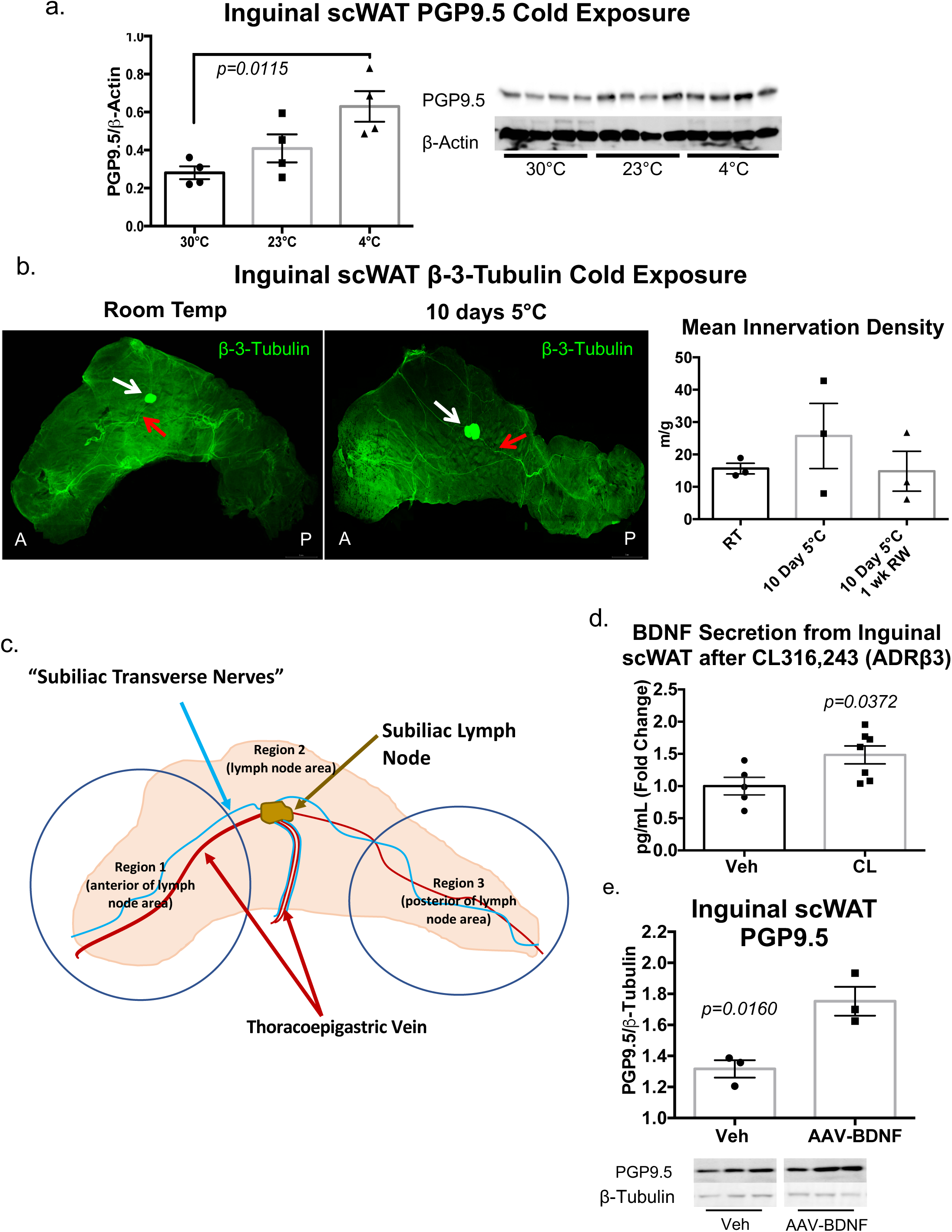
Cold exposure induces adipose nerve remodeling. Adult (8 week old) wild-type C57BL/6J male mice were either cold exposed (at 5°C), maintained at room temperature, or at thermoneutrality (30°C) for 3 days. Changes in innervation were assessed by measuring protein expression of a pan-neuronal marker (PGP9.5) in inguinal scWAT via western blotting (a). Protein expression was normalized to β-actin, band intensity was quantified in Image J and analyzed by two-tailed Student’s T-Test; N=4 per group. In a separate experiment, adult (18-22 week old) male control male mice on a mixed genetic background were maintained either at room temperature (RT), at 5°C for 10 days, or at 5°C for 10 days and then returned to RT for 1 week (rewarmed). Entire depots were immunostained with β3-Tubuilin and imaged on a Leica TCS SP8 or DMI6000 confocal microscope by tiling z-stacks through the entire depth of tissue (b). For quantification of arborization (b, right panel), tiles were individually Z-projected, background subtracted, thresholded into binary images, and skeletonized. Branches less than 4μm in length were excluded from the analysis. White arrows point to subiliac lymph node, red arrows indicate branches of the thoracoepigastric vein; anterior side (A) is left of subiliac lymph node, posterior side (P) is right of subiliac lymph node. Images are representative of N=3 per group, error bars are SEMs. Schematic of inguinal scWAT (c) divided into three anatomically distinct areas illustrating branches of the subiliac transverse nerve (blue lines) major vasculature (red lines) and centrally located subliac lymph node (gold arrow). Adult (12-13 weeks old) male C57BL/6J mice received either daily i.p. injections of ADRβ3 agonist CL316,243 (at 1.0 mg/kg BW), or vehicle (Veh) for 10-14 days; *ex vivo* secretions (collected at 1hr and 2hr) from inguinal scWAT explants were measured for BDNF by ELISA, analyzed by two-tailed Student’s T-Test, and presented as fold change in picograms (pg)/mL (d). Data is representative of multiple cohorts, N=5 (Veh), N=7 (CL316,243). Error bars are SEMs. 16 week old male BTBR MUT mice (N=3) received single injection of AAV-BDNF (1×10^10^ vg) into their left inguinal scWAT and an equal volume of vehicle into their right inguinal scWAT; protein expression of PGP9.5 in inguinal scWAT was measured by western blotting after 2 weeks (e). Protein expression was normalized to β-tubulin, band intensity was quantified in Image J and analyzed by two-tailed Student’s T-Test; error bars are SEMs.

After cold exposure, a distinct change in the neural arborization pattern was observed (Fig. 4b, middle panel), particularly an increase in intensity around the lymph node. Since these 2D representations of 3D data do not convey accurate changes in neurite density, tissue z-stacks were analyzed for mean innervation density (calculated as total axon length normalized to tissue weight) and this revealed a trend for increased innervation density in the 10 day cold exposed group compared to RT and rewarmed groups (Fig. 4b, right panel). However, taking an average of the entire tissue (as we did in Fig. 4b quantifications) blunts regional anatomical differences in innervation density, which we can observe when qualitatively assessing the tissues. This can also be seen when looking at the 2D representations in Fig. 4b, where the anterior and posterior portions display differences in innervation status after cold.

Of note, average innervation density per inguinal scWAT depot totaled up to 25 meters per tissue depot, underscoring the great density of nerve fibers contained in WAT depots. Furthermore, the inguinal scWAT depot can be considered to have 3 distinct anatomical regions, as illustrated in Fig. 4c. Region 1 is the anterior side of the depot and contains a major branch of the thoracoepigastric vein (TEV) (Fig. 4c, red arrows); the subiliac drainage lymph node, often used as an orientation landmark in inguinal scWAT is located in Region 2 as is the inferior branch of the TEV; Region 3 is the posterior end of the tissue with smaller vasculature including branches of the common iliac vein. From the anterior to the posterior side of the tissue and travelling through the subilaic lymph node is a large branching network of nerves which we have called the subliac transverse nerves (Fig. 4c, blue lines). These nerves were present in the inguinal scWAT regardless of age, exercise intervention, or cold exposure, and travelled in parallel to the main vasculature in the depot.

In separate experiments, we mimicked cold exposure by delivering the beta-3 adrenergic agonist CL316,243 (‘CL’) to wild type mice, and measured secretion of the neurotrophic factor BDNF from scWAT tissue explants. This approach revealed a significant increase in BDNF secretion after 10-14 days of CL treatments (Fig. 4d), supporting the notion that cold stimulation increases BDNF secretion from adipose, while neuropathic states reduce BDNF expression in adipose. To test whether BDNF is capable to promote scWAT innervation, we used adeno-associated virus (AAV) to deliver BDNF directly to the inguinal scWAT of BTBR MUT male mice at 16 weeks of age, when adipose neuropathy is prominent. The AS/Rec2 dual cassette vector used to deliver the BDNF transgene, transduces adipose tissue efficiently while restricting off-target transduction in liver^37^. Animals received a bolus injection of 1×10^10^ vg of AAV-BDNF in one inguinal scWAT depot and a vehicle injection in the other inguinal scWAT. After two weeks, protein expression of PGP9.5 was significantly increased in the fat pads that received virus compared to the vehicle treated depots (Fig. 4e). No difference was seen in protein expression of TH or PSD95 following AAV-BDNF treatment (Suppl. Fig. 8a-b) for this dose and duration. Taken together, these data suggest a role for BDNF in promoting total innervation on inguinal scWAT.

### Neuro-vasculature interactions in inguinal scWAT

Branches of the thoracoepigastric vein (Fig. 4b-c, red arrows) were highly innervated in all conditions and most pronounced after 10 days of cold exposure. Luxol Blue myelin staining revealed that at least some of these vascular-associated nerves are myelinated (Suppl. Fig. S9a). We had already demonstrated that nerves surround autofluorescent vasculature (Fig. 2d; 3e, Suppl. Fig. 9b), but additional assessment with the aid of the vascular fluorescent stain isolectin (IB_4_) along with PGP9.5 immunostaining, revealed further anatomical relationships between vasculature and nerves in inguinal scWAT (Suppl. Fig. S9c). Larger nerve bundles were supported by capillaries wrapping around them, while in other cases blood vessels and nerves ran parallel to one another but did not appear to directly interact. Taken together, our data present 3 distinct relationships between adipose vasculature and innervation, 1) small nerve fibers surround larger vessels, 2) vasculature envelops large nerve bundles, and 3) nerves and vessels travel together.

### Extent of adipose innervation is depot, but not sex, specific

To determine whether a difference in adipose innervation between depots and sexes exists in healthy animals, 16 week old male and female control mice were cold exposed for 3 days. No difference between sexes was observed for sympathetic activation (TH) or synapses (PSD95) in inguinal scWAT and iBAT (Suppl. Fig. S10a-b), however, both TH and PSD95 protein expression were much greater in iBAT than inguinal scWAT. Similarly, protein expression of PGP9.5 and TH in axillary and inguinal scWAT depots was comparable between sexes (Suppl. Fig. S10c-d). However, while PGP9.5 expression remained the same between the depots (Suppl. Fig. S10c), axillary scWAT exhibited significantly more protein expression of TH than inguinal scWAT (Suppl. Fig. S10d), which correlates to the ‘browning’ potential of this particular WAT depot in comparison to inguinal WAT.^38^

Since we had seen decreases in BDNF in both obesity and age-related inguinal scWAT neuropathy (Fig. 2c, Suppl. Fig. S1h), an increase in BDNF in response to noradrenergic stimulation (Fig. 4d), and observed the ability of BDNF to increase total innervation as evidenced by AAV-BDNF delivery to inguinal scWAT (Fig. 4e), we decided to investigate whether there were sex differences in BDNF expression in inguinal scWAT. After a 3-day cold exposure, adult (16 weeks old) female mice (C57BL/6J) showed a much greater level of BDNF expression in inguinal scWAT than male mice, as measured by multiplex ELISA (Suppl. Fig. S10e). This fits with the apparent protection from neuropathy in female BTBR *ob/ob* mice (Suppl. Fig. S1b and ^28^), despite females having no difference in total innervation versus males (Suppl. Fig. S10), providing the possibility that females have more adipose-secreted BDNF to maintain nerve integrity in neuropathic conditions.

## Discussion

Our data have demonstrated that under certain pathophysiological conditions, including aging, obesity and diabetes, WAT from humans and mice does not maintain proper innervation and undergoes a process of nerve death that we call ‘adipose neuropathy.’ While the exact neuropathy phenotypes differed slightly between these metabolic conditions, the underlying theme was decreased adipose tissue health. In all cases, the neuropathy was accompanied by a loss of synaptic markers and a reduction in the local expression of the neurotrophic and nerve survival factor, BDNF. These new findings have implications for the treatment of metabolic diseases, in that re-innervating adipose tissue may be required to properly regain metabolic control. Therapeutic interventions that act to support neurite outgrowth and synapse formation on adipocytes and stromovascular cells may assist with the efficacy of glucose-lowering drugs or diet and exercise interventions. In this study, we have found a promising novel treatment by locally delivering scWAT with AAV-BDNF, which appeared to succeed in re-innervating the tissues of diabetic mice.

Interestingly, while diet and exercise are often prescribed in concert, despite exercise not being a very effective means to reduce appetite, the myriad health benefits of exercise may help to bolster additional weight loss strategies. This may be in part by mediating increased peripheral nerve plasticity, including in adipose depots, as we have demonstrated here. Furthermore, we have also demonstrated that cold exposure similarly boosts peripheral nerve plasticity in adipose, and may also be a strategy to enhance diet and exercise-based weight loss interventions.

We have demonstrated that both mouse and human scWAT undergo neuropathy with aging and obesity, and these data provide evidence that peripheral neuropathy is not restricted to classical tissues such as epidermis and distal limbs. Adipose tissue nerves are known to be essential for energy-expending processes such as lipolysis and thermogenesis, and loss of a proper nerve supply can have serious detrimental effects on metabolic control, which may exacerbate or initiate an insulin resistant state. In fact, except for our BTBR mouse model and 2 individuals in each of the human cohorts (see Suppl. Table S1-S3), our adipose innervation assessments were performed under non-diabetic conditions, suggesting that adipose neuropathy may precede insulin resistance.

In all cases, lack of adipose innervation was associated with one or more of the following: an increase in adipose depot size, loss of proper adipose tissue function and metabolic control, or altered cytokine inflammatory markers. These changes in WAT may be due to loss of synapses on adipocytes themselves, or on stromovascular cells resident in WAT, including the macrophages and other immune cells that are important for proper tissue function. Indeed, macrophages are known to express adrenergic receptors^39^, and may also become denervated (neuropathic) in the models described here. Neuropathy appeared to be most severe in the obese and diabetic BTBR *ob/ob* mice, which displayed lack of synaptic integrity in the neuromuscular junction and BAT, in addition to skin and underlying adipose.

In the aged mouse model, most striking was the loss of innervation around the vasculature of WAT, without any apparent deficits to the vasculature morphology or to vascular markers such as CD31 and VEGFa (at least at the mRNA level). However, these markers cannot attest to the functionality of adipose vasculature, which could not be assessed here. Aging is known to cause alterations in structure and function of vasculature^40^ and decreased basal limb blood flow^41^, which may be a result of loss of nerve supply to the blood vessels, but to our knowledge this has not yet been investigated. Exercise was able to at least partially restore the innervation around the adipose vasculature in the aged mice. Thus, exercise may be important for prevention of aging-related diabetes by maintaining adipose innervation and metabolic health, similarly to how exercise induces neural plasticity in the brain^42^.

It is still unclear exactly how nerves and blood vessels interact and whether their plasticity is functionally linked, and this warrants further investigation. Vasculature within the adipose tissue promotes metabolic health by preventing adipose tissue hypoxia and subsequent inflammation, and provides endocrine communication between adipose tissue and the rest of the body. Nerves also rely on vasculature for sustained health, since they become damaged in a hypoxic state, while vasculature relies on innervation for constriction and vasodilation of blood vessels. Since nerves, such as those in the sympathetic nervous system, regulate vascular control (ie: vasoconstriction) and may also be involved in angiogenesis, adipose neuropathy may also have adverse effects on endocrine system communication with adipose tissue and may lead to adipose hypoxia due to lack of proper vascular supply. The association here could also go in the other direction, with circulating factors affecting nerve supply (ie: glucose, lipids, hormones) – but these hypotheses require closer investigation. The observation that nerves and blood vessels interact in three distinct ways in scWAT led us to question whether these nerves are distinct in their phenotype or function, and this is something that also warrants further exploration.

Synaptic markers and the neurotrophic factor BDNF were reduced with adipose neuropathy, fitting with the loss of pan-neuronal protein expression in neuropathic WAT depots. Neurotrophic factors, such as BDNF, are important mediators of neural plasticity, essential for nerve survival and growth, neurite outgrowth and branching, and synaptogenesis. Loss of neurotrophic signaling can prompt neurite retraction, axonal degeneration, and nerve death. It is well known that exercise often leads to an increase of BDNF in the brain ^43,44^ and in the peripheral circulation,^34^ and that this improves health outcomes in patients with neurodegenerative diseases such as Parkinson’s ^45^. It has been shown that BDNF is expressed in adipose tissue, but it’s role there is vastly understudied ^46^. Here we present a case for a potential role of BDNF in maintaining WAT peripheral nerve innervation. This idea is strengthened by the efficacy of AAV-BDNF treatments to scWAT and the striking increase in adipose BDNF in response to exercise, a known neurogenic intervention in the brain as well^42^.

Finally, we have revealed that iBAT and scWAT differ in their patterns of innervation and response to neuropathic stimuli, and that male and female mice differ in scWAT innervation with a blunted response to neuropathy and higher levels of BDNF. Using a new technique for whole-depot imaging of adipose innervation, we have also revealed a consistent pattern of nerves in inguinal WAT (traversing the subiliac lymph node), which changes regionally in response to cold stimulation.

Taken together, we have demonstrated for the first time that adipose tissue nerves are able to undergo neuropathy in pathophysiological situations, but can also undergo neural plasticity in response to exercise or cold exposure. We have also identified BDNF as a locally produced peripheral nerve survival factor in WAT, which can therapeutically improve adipose neuropathy of diabetes. Changes in adipose innervation status were correlated with altered metabolic control and underscore the importance of adipose nerves for proper metabolic health.

## Supporting information

## Acknowledgements

The authors wish to thank the following individuals from University of Maine: Brenda Kennedy-Wade and Prof. James Weber for assistance in the animal facility, Dawna Beane for paraffin embedding, Callie Greco for animal care, Jacob Cote and Mary Lander for technical assistance, Prof. Alan Rosenwasser for statistical advice, and Prof. Anne Lichtenwalner and Prof. Marie Hayes for advice. We thank Dr. Dustin Updike at MDI Biological Laboratory for microscopy assistance, and Mike Hirschman at Joslin Diabetes Center for advice on running wheel cages. We wish to thank Dr. Lei Cao at the University of Ohio for creating and gifting to us with the AAV-BDNF construct and providing advice. We also thank Dr. Ian Meng at University of New England for his kind gift of tissues from *Nav 1.8-Cre x tdTomato* reporter mice. We thank Nick Cutter, Greg Cox, and Robert Burgess for assistance with protocols for staining and quantifying neuromuscular junctions. We also wish to thank Dr. Matthew Lynes at Joslin Diabetes Center for a critical reading of the manuscript. MB was funded in part by the School of Biology and Ecology and the Graduate School of Biomedical Science and Engineering (GSBSE) at University of Maine. KLT was funded by a Junior Faculty Award from the American Diabetes Association (1-14-JF-55), a seed grant from the UMaine ADVANCE Rising Tide program, a small grant from the Boston Nutrition and Obesity Research Center (BNORC) for obtaining human adipose tissue samples, and funds from the office of the Vice President for Research at University of Maine. KLT and MB are the guarantors of the work in this manuscript.

## Author Contributions

MB created and optimized protocols, designed experiments, collected and analyzed data, and wrote the manuscript. ALD, CPJ and SW oversaw the BTBR studies, and collected and analyzed data. JWW optimized protocols, designed experiments, and conducted the majority of the microscopy studies. WPB and KBT conducted 2-photon and SHG microscopy studies. KD and LLL collected and analyzed data. MM and BH quantified the nerve fiber densities from the whole depot tiled z-stack images. KLT wrote the manuscript, designed and analyzed the experiments, and oversaw the project.

