## Supplementary material for "Neuropathy and neural plasticity in the subcutaneous white adipose depot"

*Blaszkiewicz et al. 2018*

**Supplemental Material**

Supplemental Methods

*Von Frey Analysis*

A Von Frey mechanical nociceptive assay was performed on BTBR mice ranging from 12-24 weeks of age, to determine tactile sensitivity of hind paw skin, according to protocol described by Feldman *et al.* 2009. Briefly, each mouse was subjected to five filaments (*Semmes-Weinstein* evaluators) at varying strengths (4.56, 4.31, 4.08, 3.61, 2.36), which corresponded to specific target forces (4, 2, 1, 0.4, 0.02 – grams of force (g), (Stoelting Co., Wood Dale, IL)). For the Von Frey evaluations, mice were placed on top of a grid platform into individual clear-walled compartments and allowed to acclimate with no stimulus for at least 20 minutes. After the acclimation period, trials began and filaments were applied in order of decreasing strength. Each filament was applied to the mid-plantar surface of the hind paw and slight pressure applied until the mouse showed a response or the filament bent with force. Mouse response was recorded as either positive (immediate paw removal or paw licking when filament is applied), neutral (delayed paw removal), or negative (no reaction), and each filament strength test was performed in 5 cycles.

*DRG collection*

Animals were euthanized and T13-L1 DRGs were extracted as previously described ^1^. Briefly, muscle, fat and soft tissue is cut away from the spinal column; the T13 DRG pair is readily located caudal to the most caudal ribs which were used as orienting landmarks. The spinal column was removed maintaining this orientation, cut along the midline and the spinal cord was removed in a rostral to caudal direction. Meninges, which cover the ganglia were carefully removed as well. Bilateral T13-L1 DRGs were collected and immediately frozen in liquid nitrogen, for further processing as described in *RNA Extraction and Gene Expression* section below.

*CL Injections*

Adult (12-13 week old) male C57BL/6 mice received daily i.p. injections of ADRβ3 agonist CL316,243 (Tocris Bioscience, Bristol, U.K.; Cat # 1499), at 1.0 mg/kg BW or an equivalent amount of sterile saline, for 10-14 days.

*Collection of adipose secretions and BDNF ELISA*

Inguinal scWAT depots were dissected, weighed and minced in a petri dish containing DMEM (high-glucose, serum-free). Minced tissue was transferred to a 15mL conical tube, with 5mL DMEM (loosely capped to keep tissue oxygenated) and placed in a shaking water bath at 37°C. Secretions were collected at time 0, 1hr, 2hrs, and 3hrs (1mL collected from conical tube at each time point and replaced with 1mL fresh DMEM). Secretions were stored at -80°C until processing. For ELISA, protein secretions were concentrated using Amicon^®^ Ultra Centrifugal Filters, Ultracel^®^-100K (Millipore, Burlington, MA USA; Cat. # UFC510096), per manufacturer’s instructions. Mouse BDNF PicoKine™ ELISA Kit (Boster Biological Technology, Pleasanton, CA, USA; Cat# EK0309) was used per manufacturer’s instruction to determine amount of BDNF present in adipose active secretions.

*BDNF Mulitplex ELISA*

BDNF expression in protein lysates of inguinal scWAT from 3-day cold exposed adult (16 weeks old) male and female C57BL/6J was measured using MILLIPLEX MAP Mouse Myokine Magnetic Bead Panel (Millipore, Burlington, MA USA, Cat. # MMYOMAG-74K; MILLIPLEX Magnetic Microspheres Cat. # RBDNF-MAG). Protein lysates were prepared as described under *Western Blotting* (below). The Luminex xMAP MAGPIX® system (Austin, TX, USA) was used to detect BDNF in protein samples, data was analyzed with MILLIPLEX® Analyst 5.1 software (Millipore, Burlington, MA USA) accounting for then 2-fold dilution, then normalized to total protein (previously determined by Bradford Assay).

*AAV-BDNF delivery*

16 week old male BTBR MUT mice were injected once in the left inguinal scWAT with 1x10^10^ vg of AAV-BDNF, while the right inguinal scWAT received an equal volume injection of vehicle (AAV buffer). Virus was constructed by Dr. Lei Cao as previously described ^2^. Animals, were carefully observed and scored for malaise for 48 hours after virus injection, and then observed daily, and showed no adverse reaction to the treatment. After 2 weeks animals were sacrificed, inguinal scWAT depots were harvested and processed for western blot analysis.

*Mouse adipose tissue collection and analyses; immunostaining*

Mice were euthanized, whole subcutaneous white adipose tissue (scWAT) depots were carefully removed to remain intact depots, and fixed in 2% PFA at 4°C for 4hr-12hrs depending on thickness of tissue. The tissues were then rinsed for 10 minutes with 1X PBS w/ 10u/mL heparin, twice at 4°C. Tissues were incubated in blocking buffer (1XPBS/2.5% BSA/0.5-1% Triton) at 4°C at least overnight but no more than 7 days, depending on tissue size, with blocking buffer replaced every 24hr period. After blocking period, tissues were flattened by being placed between two large glass slides bound tightly together, for at least 30min but no more than 1.5hrs at 4°C. Tissues were next incubated with 0.03% Sudan Black for 20 minutes at room temperature on a rotating platform to minimize autofluorescence. Following Sudan Black incubation, tissues were washed with 1X PBS w/ 10u/mL heparin on rotating platform at 4°C replacing PBS every 1hr for a total of 4-6hrs, or until all unbound stain was removed. Immunostaining of innervation with primary antibodies was performed overnight at 4°C, and the following day tissues were washed with 1XPBS on a rotating platform at 4°C, replacing PBS every 1hr for a total of 4-6hrs followed by incubation with secondary fluorescent antibodies.

Primary antibodies included: PGP9.5 (1:1000, Abcam, Cambridge, U.K. Cat. #10404 and #108986); post synaptic density protein 95 (1:1000-1:5000, Abcam, Cambridge, U.K. Cat. #18258); TH (1:250, Millipore, Burlington, MA USA; Cat. # AB152); NAv1.8 (1:500, StressMarq, Victoria, BC, Canada; Cat #SMC-342D); neurofilament-M (2H3, 1:500), beta-3 tubulin, (6G7, 1:250) and synaptic vesicles (SV2, 1:250) from Developmental Studies Hybridoma Bank, (University of Iowa, USA). Secondary antibodies included Alexa Fluor 488 at 1:500 and Alexa Fluor 594 at 1:1000 from Molecular Probes. For vascular autofluorescence visualization tissues were not washed in 1X PBS w/ 10u/mL heparin, prior to further immunostaining. For vascular staining, tissues were incubated with isolectin IB_4_ stain conjugated to Alexa 594 (ThermoFisher Scientific, Waltham, MA, USA; Cat # I21413) at 1μg/mL concentration overnight at room temperature. Tissues were washed in 1X PBS and mounted on slides. Images were acquired using Nikon Eclipse E400 epiflourescent microscope, Nikon A.1 confocal microscope (Nikon, Minato, Tokyo, Japan) or Leica TCS SP8 (Leica Microsystems, Wetzler, Germany) digital lightsheet/confocal microscope.

*BTBR whole depot imaging*

All 2-photon microscopy studies used a modified Olympus FV300 system with an upright BX50WI microscopy stand (Olympus, Center Valley, Pennsylvania) and a mode-locked Ti:Sapphire laser (Chameleon, Coherent, Santa Clara, California). Laser power was modulated via an electro-optic modulator (ConOptics, Danbury, Conneticut). The fluorescence and SHG signals were collected in a non-descanned geometry using a single PMT (H7422 GaAsP, Hamamatsu, Hamamastu City, Japan). Emission wavelengths were separated from excitation wavelengths using a 665 nm dichoric beamsplitter followed by 582/64 nm and 448/20 nm bandpass filters for Alexa 488 and SHG signals respectively (Semrock, Rochester, New York). Images were acquired using circular polarization with excitation power ranging from 1- 50 mW and a 40x 0.8 NA water immersion objective with 3x optical zoom with scanning speeds of 2.71s/frame. All images were 515 x 512 pixels with a field of view of 85 μm.

*Neuromuscular junction immunofluorescence, imaging, and analysis*

Following protocols provided by Greg Cox and Robert Burgess at Jackson Laboratory, mice were euthanized, both soleus and medial gastrocnemius tissues were carefully removed and fixed in a 2% PFA at 4°C for 2 hours. The tissues were then gently rinsed with 1XPBS and incubated in blocking buffer (1XPBS/2.5%BSA/0.5-1%Triton) at 4°C for at least 24 hours but no more than 7 days. After blocking period muscles were teased, tendons and fat were removed and muscle tissue was flattened by being placed between two tightly-bound glass slides for at least 30 minutes at 4°C. Tissues were next transferred to fresh blocking buffer at 4°C for at least 12 hours, but no more than 7 days. Immunostaining of innervation with primary antibodies was performed overnight at 4°C, the following day tissues were washed with 1XPBS on a rotating platform at 4°C replacing PBS every 1hr for a total of 4-6hrs. Tissues were then incubated with secondary fluorescent antibodies overnight and washed again in 1XPBS on a rotating platform at 4°C replacing PBS every 1hr for a total of 4-6hrs. Primary antibodies included: neurofilament-M (2H3, 1:500) and synaptic vesicles (SV2, 1:250) from Developmental Studies Hybridoma Bank, (University of Iowa, USA). Secondary antibodies included: Alexa Fluor 488 at 1:500 (A21121) and alpha-bungarotoxin (BTX)-conjugated to Alexa Fluor 594 at 1:1000 (B13423) from Molecular Probes (Eugene, OR, USA). Tissues were next mounted on microscope slides using Millipore mounting fluid (Burlington, MA USA; Cat. # 5013) and 1 1/5 coverslips then sealed and allowed to set overnight. All images were acquired with Nikon Eclipse E400 epiflourescent microscope using an integrated ‘Real Time Manual EDF’ acquisition tool in Nikon Elements Software (Nikon, Minato, Tokyo, Japan). Insert images were cropped in FIJI. Brightness, contrast, and sharpness were adjusted in Microsoft PowerPoint. Up to 100 NMJs were counted for each tissue and statistics were conducted in GraphPad PRISM software (La Jolla, CA, USA) using the multiple t-tests (one-per row) function.

*Luxol Fast Blue Myelin Staining of Whole Adipose*

Mice were euthanized, whole subcutaneous white adipose tissue (scWAT) depots were carefully removed to remain intact and fixed in 2% PFA at 4°C for 4hr-12hrs depending on thickness of tissue. The tissues were then rinsed for 10 minutes with 1X PBS, twice at 4°C. Tissues were stained with Luxol Blue Myelin stain kit (Abcam, Cambridge, U.K. Cat. #ab150675). Tissues were incubated in Luxol Fast Blue Solution overnight at room temperature, briefly rinsed with distilled water and differentiated in Lithium Carbonate Solution followed by 70% Alcohol Reagent until the solution ran clear and adipocytes were colorless while nerves remained blue. Tissue was then rinsed in distilled water, mounted, and imaged on a dissecting microscope.

*RNA Extraction and Gene Expression*

Total RNA was isolated from tissues using a Trizol reagent (Zymo, Irvine, CA, USA; Cat. # R2050-1-200), Bullet Blender for lysis, and Zymo Miniprep kit (Zymo, Irvine, CA, USA; Cat. # R2052). RNA yield was determined on a Nanodrop and cDNA synthesized using High Capacity Synthesis Kit (Applied Biosystems, Foster City, CA, USA; Cat# 4368813). Real-time quantitative (q)PCR was performed with SYBR Green (Bio-Rad) and CFX96 real-time PCR detection system (Bio-Rad, Hercules, CA, USA). Relative quantification analysis and fold change was performed with all values normalized to levels of a housekeeper gene (cyclophilin). Primer sequences are listed in Supplemental Table S4.

*Western Blotting*

Protein expression was measured by western blotting analysis of tissue lysates. Whole adipose depots were homogenized in RIPA buffer with protease inhibitors in a Bullet Blender, followed by Bradford Assay, and preparation of equal-concentration lysates in Laemmli buffer. 60ug was loaded per lane of a 10% polyacrylamide gel, and following gel running proteins were transferred to PVDF membranes for antibody incubations. Tyrosine hydroxylase (TH) antibody (Millipore Cat. # AB152) at 1:1000 dilution was used with both mouse and human protein lysates. Anti-PGP9.5 antibody (Abcam Cat. #ab10404 and #ab108986) was used at a 1:1000 and 1:500 dilutions respectively, were used with both mouse and human protein lysates. Anti-PSD95 antibody (Abcam Cat. #ab18258) at 1:750 dilution was used with both mouse and human protein lysates. TH, PGP9.5, and PSD95 protein expression was normalized to one of the following housekeeping proteins, all at 1:1000 dilutions: β-tubulin (Cell Signaling Technology, Danvers, MA, USA; Cat. # 2146BC), β-actin (Abcam, Cambridge, U.K.; Cat. # ab8227), cyclophilin B (Abcam, Cambridge, U.K.; Cat. #ab16045). Secondary antibody anti-rabbit HRP-linked (Cell Signaling Technology, Danvers, MA, USA; Cat. # 7074) at 1:3000 dilution was used for conjugation with all primaries. Blots were visualized on a Syngene G:BOX (Frederick, MD, USA). Ponceau S staining was performed after immunoblotting was complete using Ponceau S Solution (Sigma-Aldrich, St. Louis, MO, USA; Cat. # P7170-1L).

*Cold Exposure Experiments*

All cold exposure was carried out in a diurnal incubator (Caron, Marietta, OH, USA) at 5°C. Animals were housed two to a cage and continuously cold exposed for 3 - 14 days.

For whole adipose innervation imaging, 18-22 week old control male mice on a mixed genetic background were housed either at room temperature, cold exposed for 10 days, or cold exposed for 10 days and returned to room temperature for 1 week (‘rewarmed’).

*Adipose clearing to visualize sympathetic innervation*

Inguinal scWAT depots from *TH-Cre x ROSA YFP* (Jackson Laboratory, Bar Harbor, ME, USA; Stock # 008601 and 006148) reporter mice were cleared using ScaleView A2 as described by Hama, H. et al. ^3^ and a sucrose gradient method as described by Tsai, P. et al. ^4^. Images were acquired on multiple platforms including Syngene G:BOX (Frederick, MD, USA), Leica TCS SP8 (Leica Microsystems, Wetzler, Germany), and Nikon A.1 (Nikon, Minato, Tokyo, Japan).

*Nav1.8 reporter mice*

Adipose tissues from young adult female *Nav 1.8-Cre x tdTomato* reporter mice (gifted by Dr. Ian Meng, University of New England) were used for whole depot immunostaining with PGP9.5 (GFP) as described in ‘*Mouse adipose tissue collection and analyses; immunostaining’* section above.

**Supplemental Figures:**

**Supplemental Figure S1: Adiposity & Neuropathy of BTBR *ob/ob* mutant (MUT) mice.** Male and female BTBR MUT mice were assessed for total body weight and adiposity (inguinal scWAT/body weight), and compared to WT or HET animals in a pilot cohort (a). Female BTBR WT, HET, and MUT were assessed for tactile allodynia via the Von Frey assay, an indirect measure of peripheral neuropathy (b). Protein expression of PGP9.5 (c) and TH (d) in inguinal scWAT of HET, MUT, and WT BTBR mice was measured via western blotting. For (a-d), males: MUT N=2; WT N=3; HET N=3; females: MUT: N=1; WT: N=3; HET: N=3; all mice were 12-24 weeks old. Data represents a pilot cohort to compare HET to MUT mice, and males to females, thus statistical analyses were not performed due to small sample sizes. Sample Ponceau S staining to demonstrate equal protein loading (e). Whole BAT depots were compared between female and male BTBR HET, MUT, and WT mice (f). Images are representative, males: MUT N=2; WT N=3; HET N=3; females: MUT N=1; WT N=3; HET N=3; all mice were 12-24 weeks old. Gene expression analysis of BAT, inguinal scWAT, and prWAT (g-i). Gene expression data were analyzed by two-tailed Student’s t-test, using Welch’s correction when variance was unequal, N=4-5 per group. Error bars are SEMs.

**Supplemental Figure S2: Adipose neuropathy of BTBR *ob/ob* mutant (MUT) mice.**

Neuromuscular Junction (NMJ) Analysis (a-b)**.** Immunofluorescent staining of male BTBR WT and MUT neuromuscular junctions of the medial gastrocnemius (MG) and soleus (SOL) muscles was performed using neurofilament M (2H3) and synaptic vesicles (SV2) (in green) to visualize the pre-synaptic area, and α-bungarotoxin (in red) to visualize the post-synaptic area. Representative images at 40x magnification of BTBR *ob/ob* wild-type (WT) medial gastrocnemius (MG) (top left panel), and soleus (bottom left panel). Mutant (MUT) MG (top right panel) and SOL (bottom right panel) (a). Inserts are of occupied (left panels) and partially occupied (partially occupied) NMJs in representative images (a). Percent of total NMJs for WT and MUT animals in both MG and SOL muscles was calculated as an indicator of neuropathic state (b). Analysis shows multiple t-tests (per row) of replicate cohorts; N=7 for WT MG, N=6 for MUT MG, N=7 for WT SOL, and N=4 for MUT SOL. Error bars are SEMs. *p < 0.05, **p < 0.01, ***p < 0.001, ****p < 0.0001. Two-photon microscopy was performed on immunofluorescent stained (PGP9.5) inguinal scWAT of WT and MUT animals (c). PGP9.5 was detected by excitation of AlexaFluor 488 at 800nm and emission collected using a 582 +/- 64nm filter. For detection of collagen, samples were excited at 890nm and the SHG signal was collected using a 448 +/-20nm filter. A 40x water immersion objective was used. IMARIS software was used to render 3D projections from z-stacks. Images are representative of N=3 WT/MUT, 12 week old males.

**Supplemental Figure S3. Human scWAT cell size and omental adipose innervation.** Average cell diameter in scWAT of human samples when assessed by BMI or age (a), and normalized protein expression plotted against average cell size for either BMI or age cohorts (b), corresponds to western blot data in Fig. 1g-h. Cell diameter measured from images of histological cross-sections of adipose tissue from each patient, averaged (n=3), and analyzed by linear regression. Human omental adipose samples were analyzed for protein expression of PGP9.5 via western blotting (c-d). Linear regression was performed with respect to BMI (c) or age (d). Protein expression was normalized to β-Actin and band intensities were quantified in Image J. Due to limitations of available samples from the BNORC adipose tissue core at Boston Medical Center, the BMI distribution of omental adipose was clustered around 40. Error bars are SEMs.

**Supplemental Figure S4: Body weight and inguinal scWAT innervation of young/aged and sedentary/exercised (run) C57BL/6J mice.** Body weight, adiposity (inguinal scWAT/body weight & pgWAT/body weight), and quadricep muscle weight was measured for young (12 weeks old) and aged (16 month old) mice under sedentary (sed) and exercised (run) conditions (a). Young and aged sedentary (sed) groups, N=4; young and aged exercised (run) groups N=5. Body and tissue weight analyzed by one-way ANOVA, with Tukey post hoc, groups labeled with the same letter (A or B) are statistically similar, for body weight: *p=0.0473* for young sed v. aged sed; *p=0.*0240 for young run v aged run. Protein expression PGP9.5 (b) and TH (c) in inguinal scWAT of young run versus aged run mice was determined by western blotting. Protein expression was normalized to β-Actin or Cyclophilin B; band density was quantified in Image J and analyzed by two-tailed Student’s t-test. Protein expression of PGP9.5 (d), TH (e), and PSD95 (f) in BAT of young (12 week old) sedentary (young sed) versus young exercised (young run) mice was determined by western blotting. Protein expression was normalized to β-tubulin or cyclophilin B; band density was quantified in Image J and analyzed using a two-tailed Student’s t-test. Gene expression analysis of BAT from young (12-15 week old) sedentary (young sed) versus young exercised (young run) male mice (g), gene expression was analyzed by two-tailed Student’s t-test, N=4 for sedentary and N=6 for run. Error bars are SEMs.

**Supplemental Figure S5: Innervation of vasculature in young sedentary/exercised (run) mice.** Whole inguinal scWAT depots were collected from young *C57BL6/J* mice under sedentary and exercised (run) conditions. Tissue was stained with PGP9.5 (green) and Isolectin Ib-4 (red) to analyze nerve and blood vessel interactions in WAT with exercise. Tissues were scanned at 10x for blood vessels 50um or greater in diameter. Twenty blood vessels were evaluated for innervation in each tissue (or as close to 20 as could be found) (a-b). Percentage of innervated vessels with a diameter of ≥ 50um were evaluated per tissue (a, left panel). Analyzed with two-tailed Student’s T-Test. Error bars are SEMs. Innervation was further characterized by the number of resident nerves per blood vessel (a, right panel). Data was analyzed using a two-way ANOVA and Tukey’s multiple comparison test. Error bars are SEMs. Fluorescent imaging of blood vessel and nerve interactions at 4x, 10x, and 40x using an E400 epifluorescent microscope (b). N=4 for sedentary and N=5 for run groups.

**Supplemental Figure S6: Adipose nerve imaging techniques.** Paraffin embedded 7um thick section of inguinal scWAT from male *TH-Cre x ROSA YFP* mice and stained with PGP9.5 imaged on epifluorescent microscope with a 40x objective (a). ScaleView-A2 cleared inguinal scWAT of 7 day cold exposed (5ºC) male *TH-Cre x ROSA YFP* mice, imaged on a Syngene G:BOX (b). Red arrows indicate changes in nerve intensity with re-warming. Whole inguinal scWAT depot from male C57BL/6J mice, collected after 14-day cold exposure (5ºC) sucrose cleared, immunostained with β3-Tubulin, imaged on a confocal microscope and presented as a depth coded tiled mosaic including a scale bar on left indicating nerve depth (c). Immunofluorescent staining of PGP9.5 (green) in whole inguinal scWAT depots from male C57BL/6J mice were imaged on an epifluorescent microscope at 4x and 10x objectives showing patterns of innervation (d). Whole depot pgWAT from C57BL/6J, was co-stained with synaptic marker PSD95 (red); and neurofilament M (2H3) and synaptic vesicles (SV2) (green), and imaged on an epifluorescent microscope with a 40x objective to show location of synapses (e). Inguinal scWAT depots taken from male C57BL/6J mice were either washed on a rotating shaker at room temperature in 1X PBS prior to staining (f, left panel) or washed in 1X PBS w/ 10u/mL heparin (f, right panel) as a means to reduce vascular autofluorescence. Tissues were then stained with PGP9.5 (green) and imaged on an epifluorescent microscope with a 10x objective.

**Supplemental Figure S7. Neuronal reporters for specific nerve subtypes compared to the respective antibodies.** Inguinal scWAT depots of female *Na_v_1.8-Cre x tdTomato* (red) mice were immunostained with PGP9.5 (green) (a, left panel) and compared to female *C57BL/6J* mice scWAT depots co-stained with Na_v_1.8 (red) and PGP9.5 (green) antibodies, imaged on an upright epifluorescent microscope with 10x objective (a, right panel). Inguinal scWAT depots of male *TH-Cre x ROSA YFP* (green) mice, imaged on confocal microscope with 5x objective, and digitally zoomed (b, left panel) and male C57BL/6J mice stained with TH antibody (red) on a confocal microscope with 10x objective (b, right panel).

**Supplemental Figure S8 – AAV mediated BDNF delivery to inguinal scWAT of BTBR MUT mice.** Adult (16 week old) male BTBR MUT mice (N=3) received single injection of AAV-BDNF (1x10^10^ vg) into their left inguinal scWAT and an equal volume of vehicle into their right inguinal scWAT. Two weeks post-injection, protein expression of TH (a) and PSD95 (b) in inguinal scWAT was measured by western blotting. Protein expression was normalized to cyclophilin or β-tubulin, band intensity was quantified in Image J and analyzed by two-tailed Student’s T-Test. Error bars are SEMs.

**Supplemental Figure S9: Neurovascular interactions**

Luxol Fast Blue myelin staining of whole inguinal scWAT depot (a). Image taken on a dissecting microscope and is a digital zoom of 30x original magnification, arrows point to branches of the thoracoepigastric vein. Immunofluorescent staining of inguinal scWAT depots from male C57BL/6J mice with synaptic vesicles (SV2) and neurofilament M (2H3) (green) and autofluorescence of vasculature (orange). Images were taken on an epifluorescent microscope at 10x magnification (b). Inguinal scWAT depots collected from male C57BL/6J mice were immunostained with PGP9.5 (green) and co-stained with Isolectin (red) to further visualize neurovascular interactions. Imaged on both confocal microscope with a 10x objective, (tiled, max projection) (c, top panels), and epifluorescent microscope with 10x and 4x objectives (c, middle and bottom panels respectively).

**Supplemental Figure S10: Adipose innervation: Sex and depot comparison**

Adult (16 week old) male and female control mice on a C57BL/6J background were cold exposed (5°C) for 3 days. Protein expression of TH (a) and PDS95 (b) were measured in inguinal scWAT and BAT via western blotting. Protein expression of PGP9.5 (c) and pan-neuronal marker TH (d) was measured in axillary and inguinal scWAT for both sexes via western blotting. Protein expression was normalized to β-Tubulin or Cyclophilin B, band density was quantified in Image J and analyzed using a two-tailed Student’s t-test. Error bars are SEMs. BDNF expression in inguinal scWAT was measured for both sexes via multiplex ELISA assay (e).

**Supplemental Materials References**

1.	Cao, Y., Wang, H. & Zeng, W. Whole-tissue 3D imaging reveals intra-adipose sympathetic plasticity regulated by NGF-TrkA signal in cold-induced beiging. *Protein Cell* **9**, 527-539 (2018).

2.	Huang, W., Liu, X., Queen, N.J. & Cao, L. Targeting Visceral Fat by Intraperitoneal Delivery of Novel AAV Serotype Vector Restricting Off-Target Transduction in Liver. *Mol Ther Methods Clin Dev* **6**, 68-78 (2017).

3.	Hama, H.*, et al.* Scale: a chemical approach for fluorescence imaging and reconstruction of transparent mouse brain. *Nat Neurosci* **14**, 1481-1488 (2011).

4.	Tsai, P.S.*, et al.* Correlations of neuronal and microvascular densities in murine cortex revealed by direct counting and colocalization of nuclei and vessels. *J Neurosci* **29**, 14553-14570 (2009).
