## Supplementary material for "Neuropathy and neural plasticity in the subcutaneous white adipose depot"

**Supplemental Table S1 - Human Cohort Data by Age^[1]^**

| **Used in**  **Figure #** | **Age** | **BMI (kg/m^2)^** | **Plasma metabolites** | | | | | |
| --- | --- | --- | --- | --- | --- | --- | --- | --- |
|  |  |  | **BG Rand (mg/dL)** | **Cholesterol (mg/dL)** | **Triglycerides**  **(mg/dL)** | **HDL**  **(mg/dL)** | **LDL**  **(mg/dL)** | **HgBA1C** |
| 1h, S2a-c | 25 | 40 | no data | 170 | 158 | 53 | 85 | 5.2 |
| 1h, S2c (age) | 27 | 29 | no data | 192 | 44 | 33 | 93 | 5.7 |
| S2a-b | 28 | 40.78 | 96 | 154 | 158 | 51 | 71 | 5.1 |
| 1h, S2c (age) | 30 | 21.69 | no data | no data | no data | no data | no data | no data |
| S2a-b | 31 | 76.88 | 97 | 179 | 93 | 41 | 119 | 5.5 |
| 1h, S2c (age) | 32 | 27.55 | 95 | 185 | 108 | 54 | 109 | 5.3 |
| 1g-h, S2a-c | 33 | 35.17 | no data | 200 | 145 | 41 | 130 | no data |
| 1h, S2c (age) | 34 | 24 | no data | no data | no data | no data | no data | no data |
| 1g-h, S2c | 34 | 40.4 | no data | no data | no data | no data | no data | no data |
| 1h | 35 | 26.82 | no data | no data | no data | no data | no data | no data |
| 1h, S2c (age) | 38 | 25 | 87 | no data | no data | no data | no data | no data |
| S2a-b | 38 | 37.7 | 98 | 175 | 132 | 39 | 110 | 6.3 |
| 1g, S2c (BMI) | 40 | 30.7 | 82 | no data | no data | no data | no data | no data |
| 1g, S2a-b, c (BMI) | 41 | 42 | no data | no data | no data | no data | no data | no data |
| S2a-b | 42 | 46.32 | 109 | 147 | 155 | 40 | 76 | 6 |
| 1g, S2c (BMI) | 47 | 30.64 | 93 | no data | no data | no data | no data | 4.9 |
| 1g, S2c (BMI) | 47 | 45.74 | 64 | 169 | 191 | 44 | 87 | 4.8 |
| 1g, S2c (BMI) | 51 | 30 | no data | no data | no data | no data | no data | no data |
| 1g, S2c (BMI) | 60 | 38.38 | 93 | no data | no data | no data | no data | no data |
| 1g, S2c (BMI) | 63 | 36 | no data | 200 | 92 | 62 | 120 | 6.7 |

1. Metabolic data from patients whose adipose tissues were assessed for innervation, data sorted by ascending age BMI = body mass index; BG Rand = random blood glucose; HDL = high density lipoprotein; LDL = low density lipoprotein; HgBA1c = hemoglobin A1c.
