## Supplementary material for "Neuropathy and neural plasticity in the subcutaneous white adipose depot"

**Supplemental Table S2 - Human Characteristic Data^[1]^**

| **Used in Figure #** | **Tissue** | **Sex** | **Procedure** | **Race/ Ethnicity** | **Diabetes** |
| --- | --- | --- | --- | --- | --- |
| 1h, S2c (age) | sqWAT | F | Panniculectomy | Hispanic | Non-diabetic |
| 1g-h, S2c | sqWAT | F | Panniculectomy | Hispanic | Non-diabetic |
| 1h, S2c (age) | sqWAT | F | Panniculectomy | African American | Diabetic |
| 1h, S2c (age) | sqWAT | F | Panniculectomy | Hispanic | Non-diabetic |
| 1g, S2c (BMI) | sqWAT | F | Panniculectomy | Hispanic | Non-diabetic |
| 1g, S2c (BMI) | sqWAT | F | Panniculectomy | African American | Non-diabetic |
| 1h, S2c (age) | sqWAT | F | Panniculectomy | Unknown | Pre-diabetic |
| 1g, S2c (BMI) | sqWAT | F | Panniculectomy | Hispanic | Non-diabetic |
| 1g, S2c (BMI) | sqWAT | F | Abdominoplasty | Unknown | Non-diabetic |
| 1h, S2c (age) | sqWAT | F | Abdominoplasty | Hispanic | Non-diabetic |
| 1h | sqWAT | F | Panniculectomy | African American | Diabetic |
| 1g, S2c (BMI) | sqWAT | F | Panniculectomy | Hispanic | Diabetic |
| 1g, S2c (BMI) | sqWAT | F | Panniculectomy | African American | Diabetic |
| 1g, S2a-b, c (BMI) | sqWat, Omental | F | Gastric Bypass | African American | Non-diabetic |
| 1g-h, S2a-c | sqWat, Omental | F | Gastric Bypass | Unknown | Non-diabetic |
| 1h, S2a-c | sqWat, Omental | F | Gastric Bypass | Hispanic | Non-diabetic |
| S2a-b | Omental | F | Gastric Bypass | Hispanic | Non-diabetic |
| S2a-b | Omental | M | Gastrectomy Sleeve | Hispanic | Non-diabetic |
| S2a-b | Omental | F | Gastric Bypass | Hispanic | Pre-diabetic |
| S2a-b | Omental | F | Gastric Bypass | Hispanic | Non-diabetic |

1. Patient tissue type analyzed, sex, elective surgical procedure, race or ethnicity, and diabetic state are presented along with which figure samples were used in. sqWAT = subcutaneous white adipose tissue; BMI = body mass index; F = female; M = male.
