## Supplementary material for "Neuropathy and neural plasticity in the subcutaneous white adipose depot"

**Supplemental Table S3 – Human Cohort Data Total Statistics^[1]^**

| **Percent of Total** | | |
| --- | --- | --- |
| **Gender** | Male | 5 |
|  | Female | 95 |
| **Ethnicity/Race** | Hispanic | 60 |
|  | African American | 25 |
|  | White | 0 |
|  | Unknown | 15 |
| **Age** | 20-29 | 15 |
|  | 30-39 | 45 |
|  | 40-49 | 25 |
|  | 50-59 | 5 |
|  | 60+ | 10 |
| **Menopausal** | Premenopausal | 70 |
|  | Postmenopausal | 1 |
|  | Unknown | 15 |
| **Diabetes** | Pre-diabetic | 10 |
|  | Diabetic | 20 |
|  | Nondiabetic | 70 |
| **BMI** | Normal: <25.0 | 10 |
|  | Overweight: 25.0-29.9 | 20 |
| **Obese** | Class I: 30.0-34.9 | 15 |
| **Obese** | Class II: 35.0-39.9 | 20 |
| **Obese** | Class III: >40.0 | 35 |
| **Procedure** | Panniculectomy | 55 |
|  | Gastric Bypass | 30 |
|  | Abdominoplasty | 10 |
|  | Gastrectomy Sleeve | 5 |

1. Summarized statistics are shown for all patients in the study. Numbers represented as percent of total.
