## Supplementary material for "Neuropathy and neural plasticity in the subcutaneous white adipose depot"

**Supplemental Table S4**

| **qPCR Primers** | | |
| --- | --- | --- |
| ***Gene*** | **Forward Sequence** | **Reverse Sequence** |
| *bdnf* | CAGGTGAGAAGAGTGATGACC | ATTCACGCTCTCCAGAGTCCC |
| *cd31* | ACGCTGGTGCTCTATGCAAG | TCAGTTGCTGCCCATTCATCA |
| *cidea* | ATCACAACTGGCCTGGTTACG | TACTACCCGGTGTCCATTTCT |
| *cgrp* | CCTGCAACACTGCCACCTGCG | GAAGGCTTCAGAGCCCACATTG |
| *cyclophilin* | CAAATGCTGGACCAAACACAA | AAGACCACATGCTTGCCAT |
| *dio2* | CAGTGTGGTGCACTGCTCCAATC | TGAACCAAAGTTGACCACCAG |
| *il4* | GGTCAACCCCCAGCTAGT | GCCGATGATCTCTCTCAAGTGAT |
| *il10* | CTATGCTGCCTGCTCTTACTGAC | CGGAGAGAGGTACAAACGAGG |
| *il13* | CCTGGCTCTTGCTTGCCTT | GGTCTTGTGTGATGTTGCTCA |
| *nyp* | AAGCCGGACAATCCGGGCCGAGG | GCTTTCCTCATTAAGAGGTCT |
| *pacap* | TACTGTGTGTGTAACTGTGTGGG | GCCAGCCGTAAGTAGATGCTC |
| *penk* | TTCAGCAGATCGGAGGAGTTG | AGAAGCGAACGGAGGAGAGAT |
| *pgc1α* | CCCTGCCATTGTTAAGACC | TGCTGCTGTTCCTGTTTTC |
| *pomc* | AGA CCT CCA TAG ATG TGT GGA | AGC GGA AGT GAC CCA TGA CGT |
| *psd95* | GCGGTGCTAAAATCGAATGC | ACAGAGAGGGGCAGGCAGT |
| *sox10* | AGATCCAGTTCCGTGTCAATAA | GCGAGAAGAAGGCTAGGTG |
| *synapsin I* | CATGGCACGTAATGGAGACTACCGCA | CCGCCAGCATGCCTTC |
| *synapsin II* | GCCACCAGGTTAAGCTCTGA | TTCCAGGAAGGCCAAGGT |
| *synaptophysin* | TGACTTCAGGACTCAACACCTC | CAGGAGCTGGTTGCTTTTCT |
| *tac1* | TGGCCAGATCTCTCACAAAAGGCA | TCGTAGTTCTGCATCGCGCTTCT |
| *ucp1* | AGGCTTCCAGTACCATTAGGT | CTGAGTGAGGCAAAGCTGATTT |
| *vegfa* | AACAAAGCCAGAAAATCACTGTGA | CGGATCTTGGACAAACAAATGC |
| *vip* | AGTGTGCTGTTCTCTCAGTCG | GCCATTTTCTGCTAAGGGATTCT |
