## Supplementary material for "Neuropathy and neural plasticity in the subcutaneous white adipose depot"

### Supplemental Data

#### Supplemental Figure S1 – Adiposity & neuropathy of BTBR *ob/ob* mutant (MUT) mice.

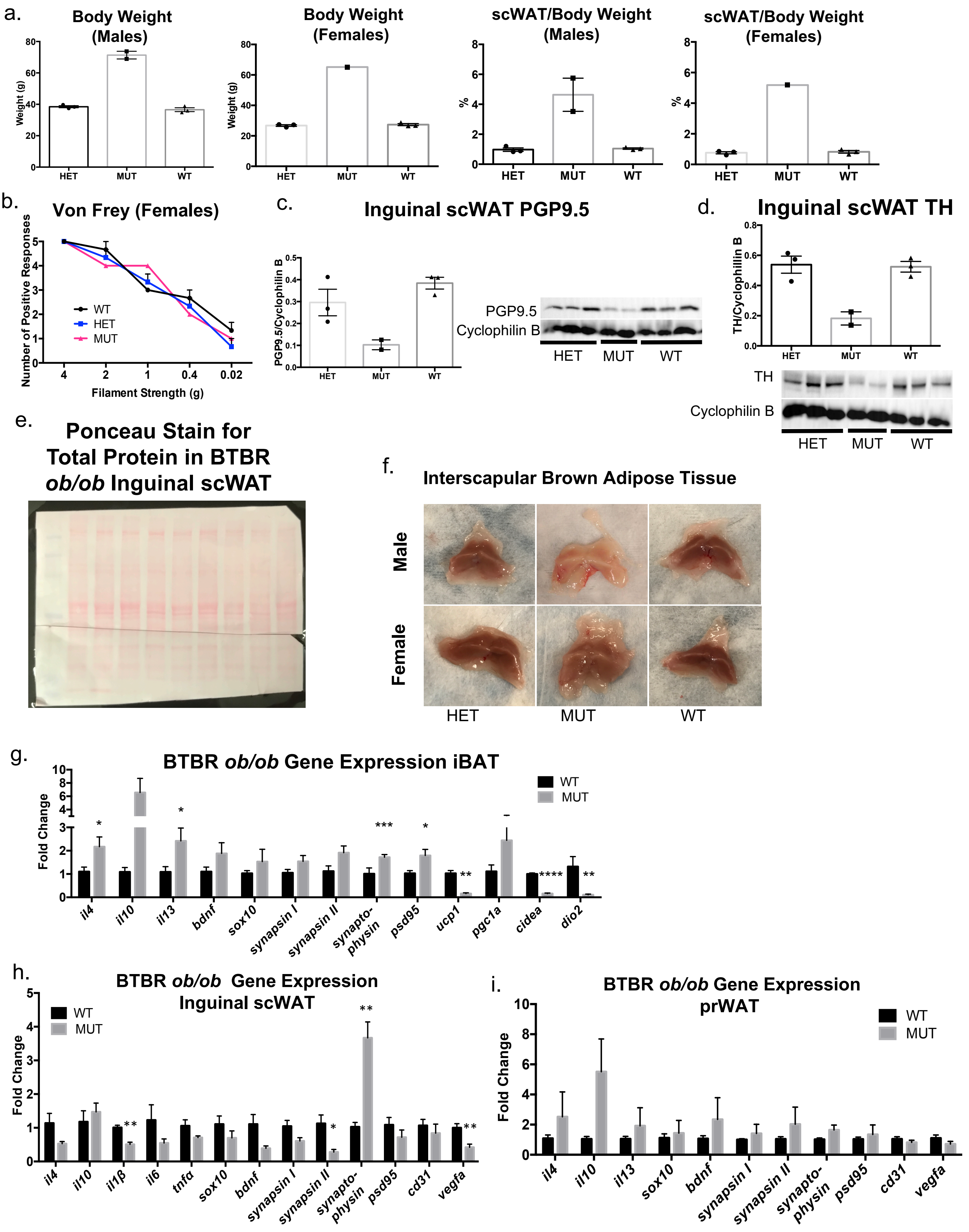

#### Supplemental Data

##### Supplemental Figure S2 – Adipose neuropathy of BTBR *ob/ob* mutant (MUT) mice.

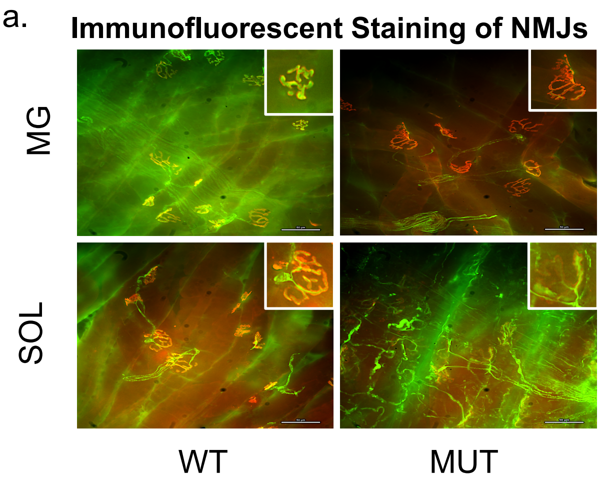


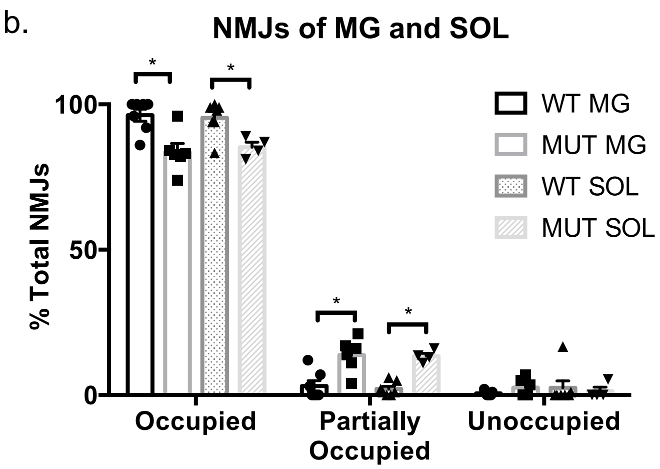


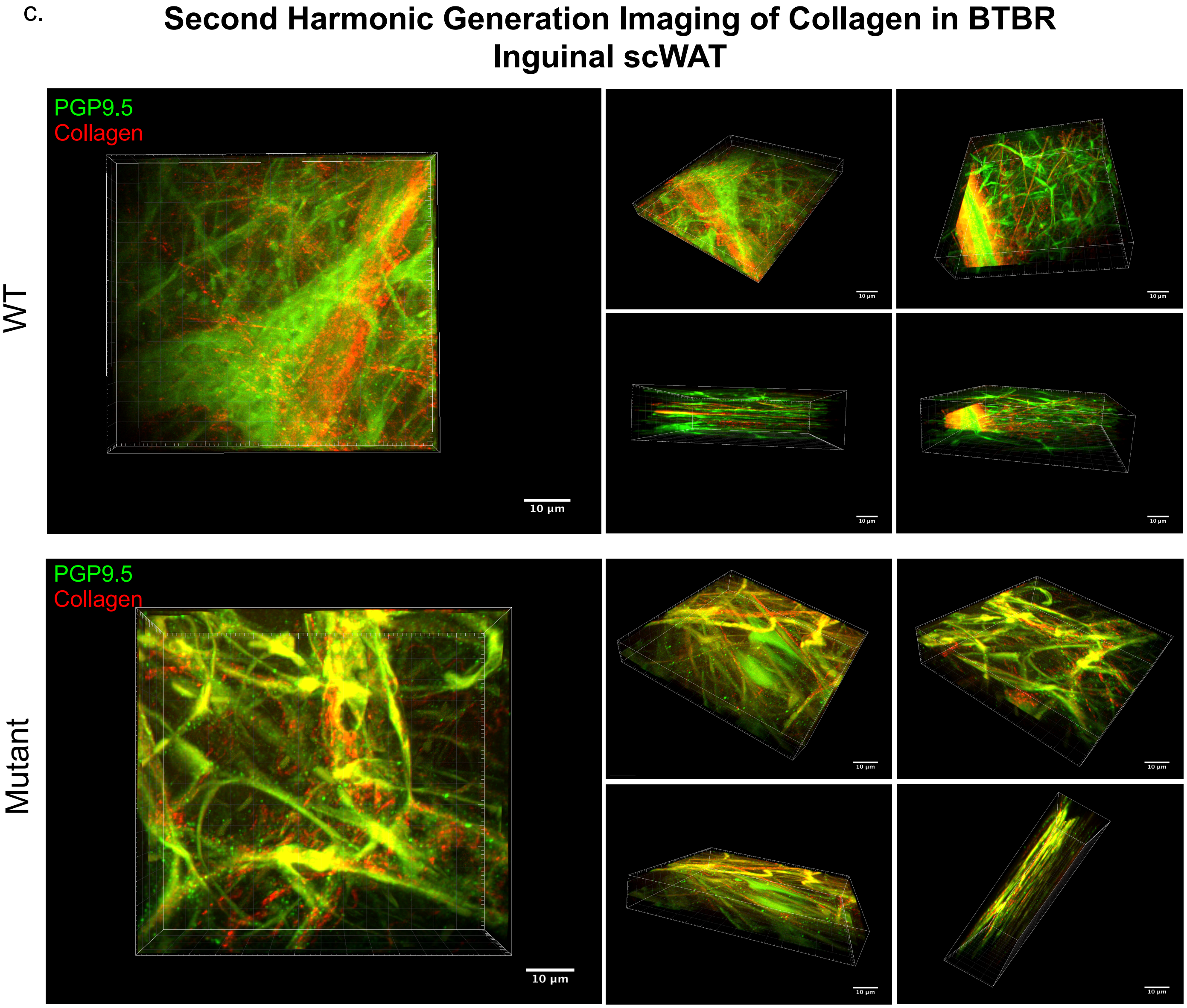

**Supplemental Data**

**Supplemental Figure S3 – Human scWAT cell size and omental adipose innervation**

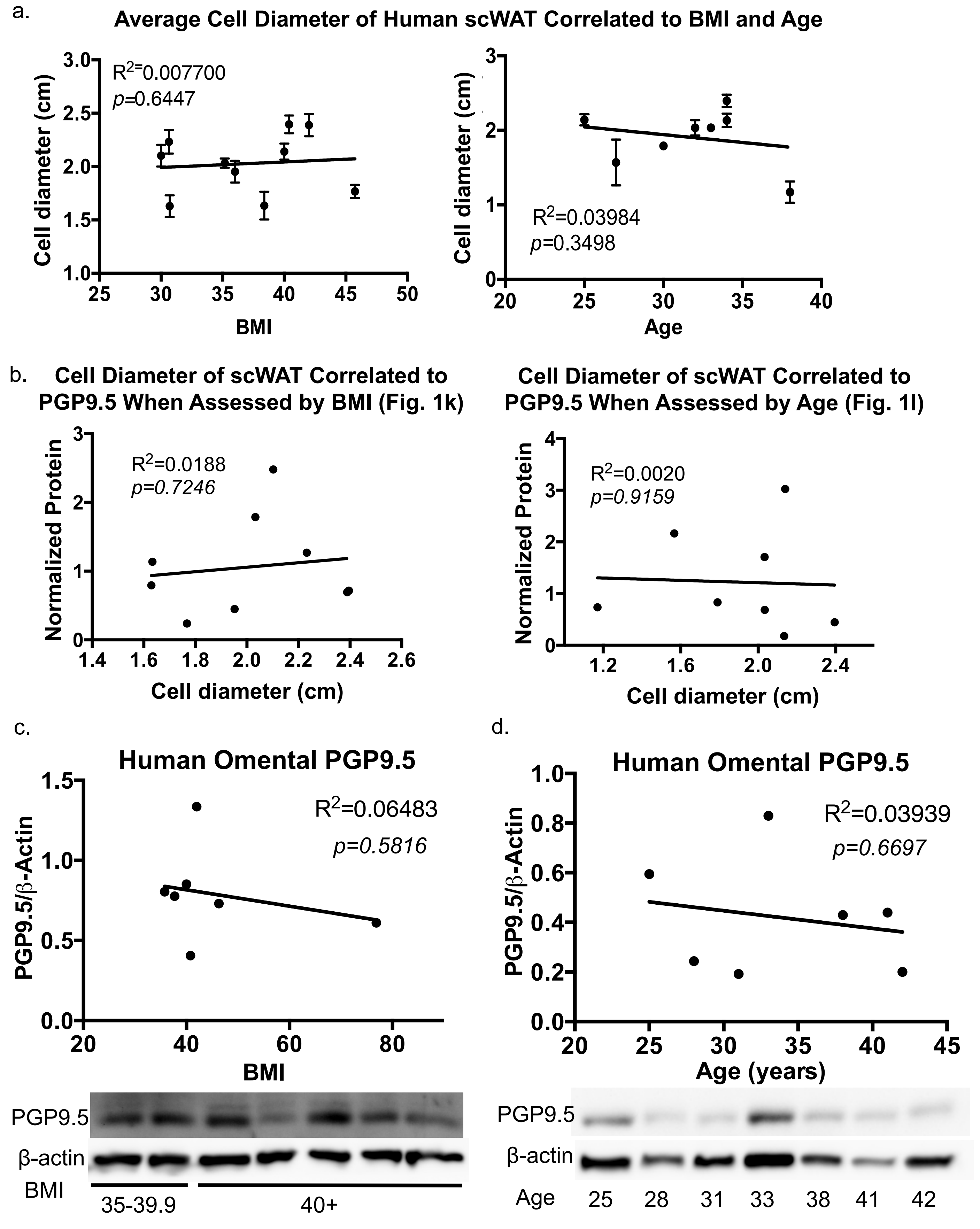

#### Supplemental Data

#### Supplemental Figure S4 – Body weight, inguinal scWAT, and BAT innervation of young/aged and sedentary/exercised (Run) mice

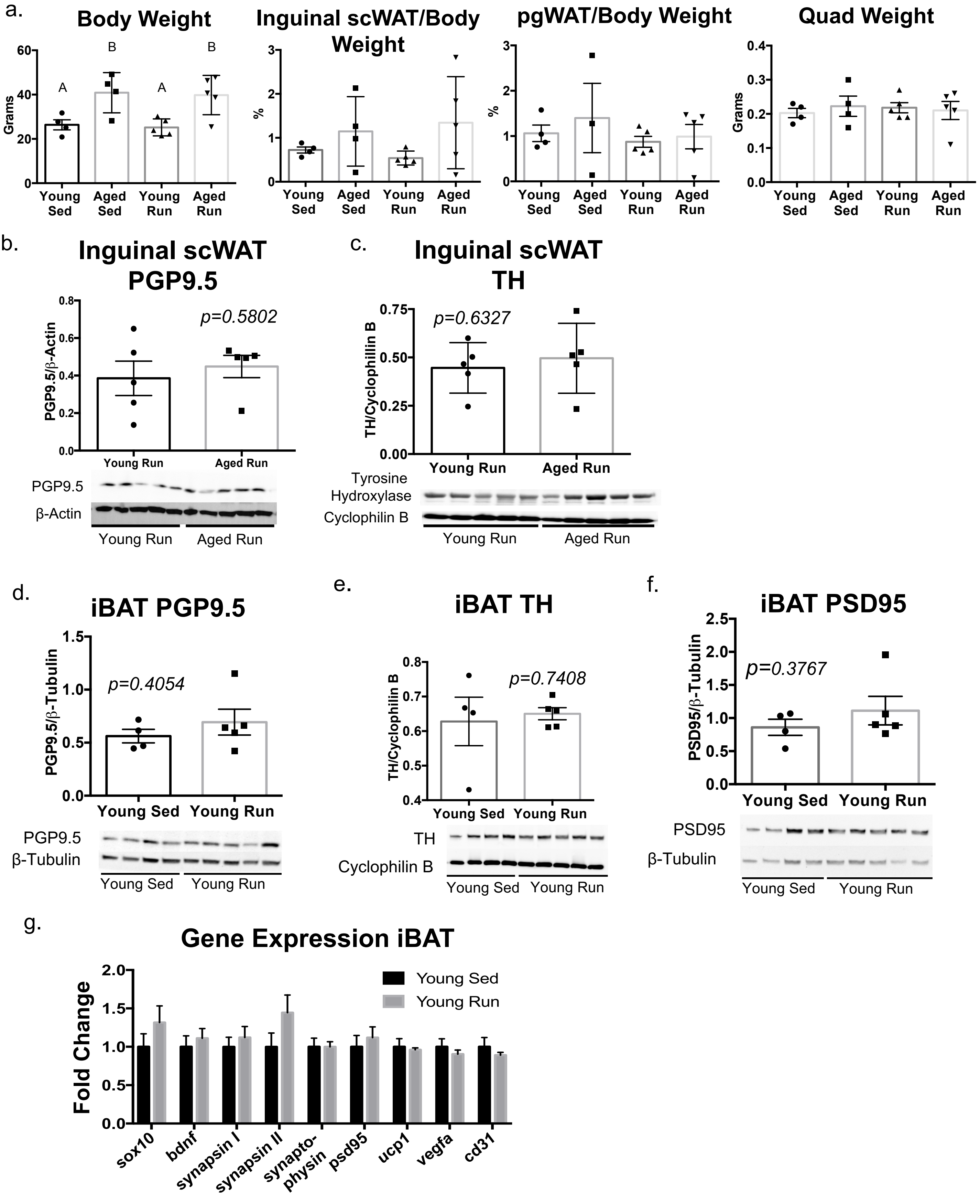

**Supplemental Data**

**Supplemental Figure S5 – Innervation of vasculature in young sedentary/exercised (Run) mice**

a.

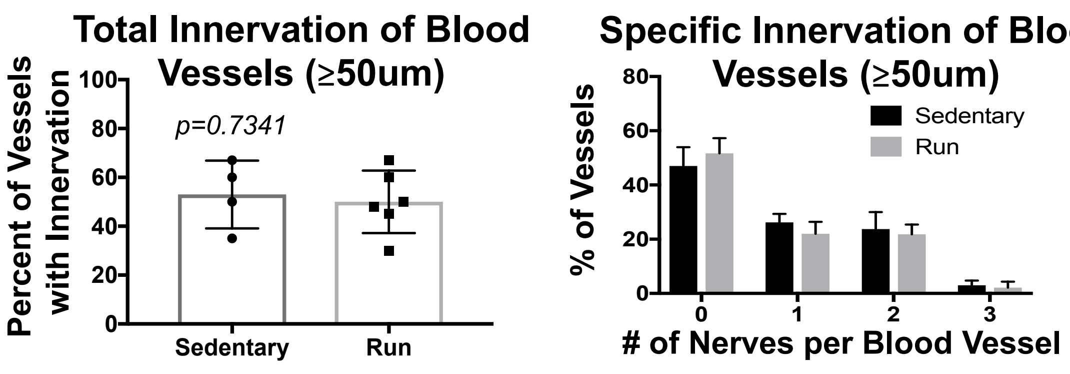


b.

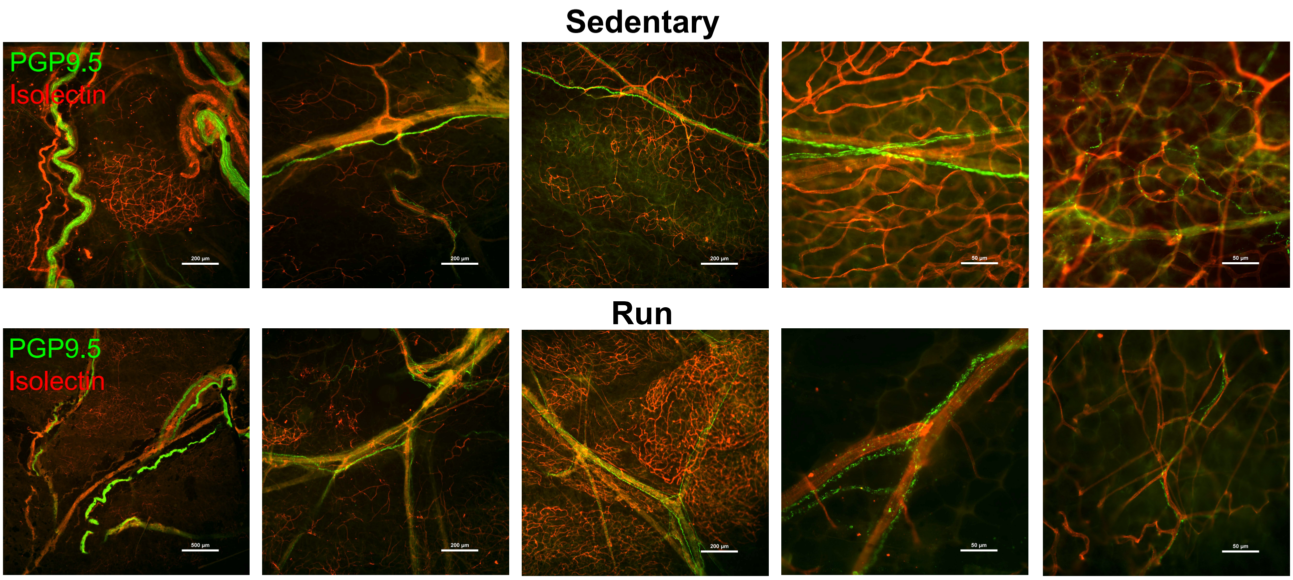

**Supplemental Data**

#### Supplemental Figure S6 – Adipose nerve imaging techniques

a. **Standard Immunohistochemistry of 7uM tissue sections**

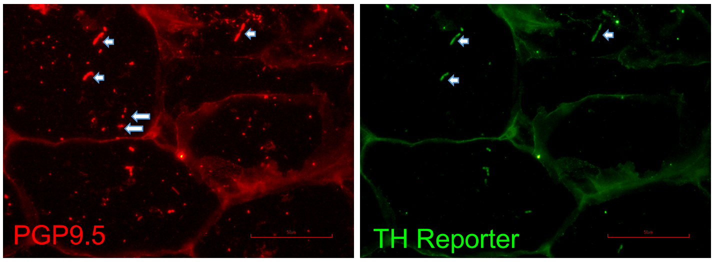


b. **ScaleView-A2 Cleared Inguinal scWAT – TH-Cre x ROSA YFP reporter**

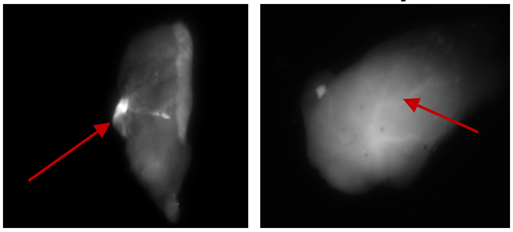

Cold    Re-Warmed

c. **Depth Coding of Inguinal scWAT Innervation after Cold Exposure**

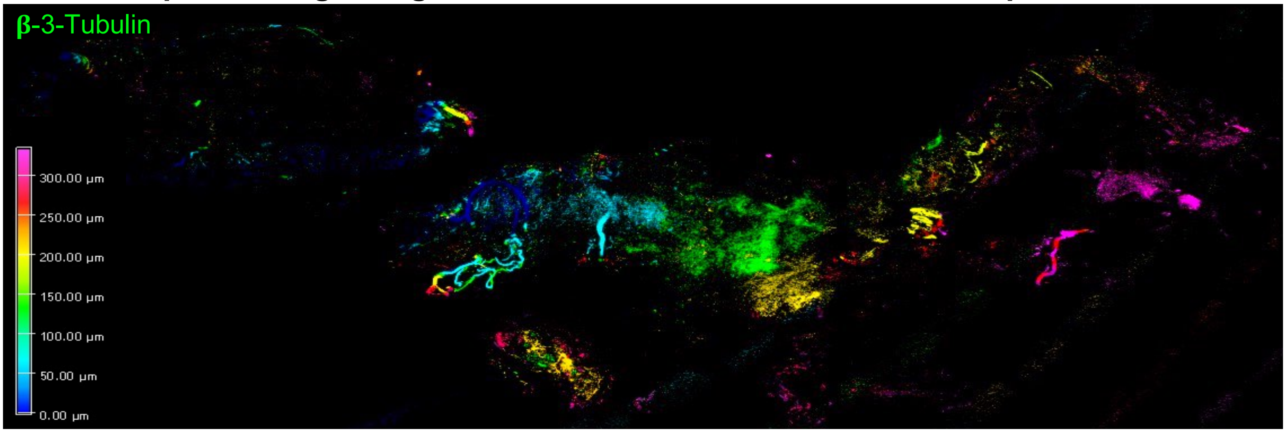


d. **Inguinal scWAT Immunofluorescent Staining**

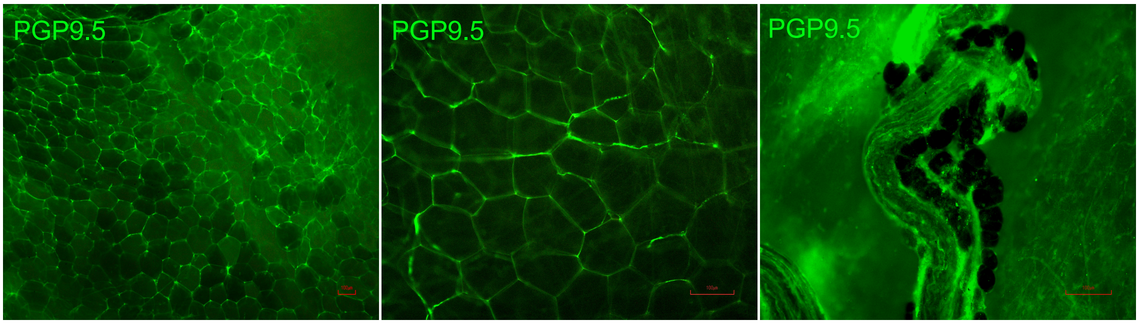


e. **pgWAT Synapses in SVF**

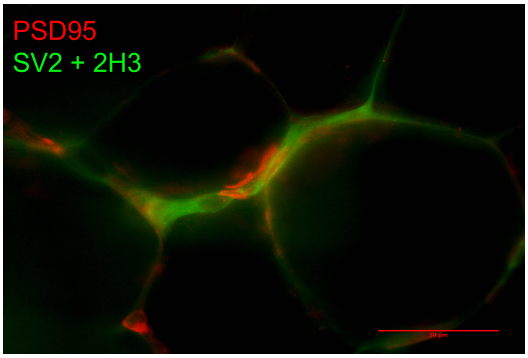


f. **Inguinal scWAT Immunostaining Optimization**

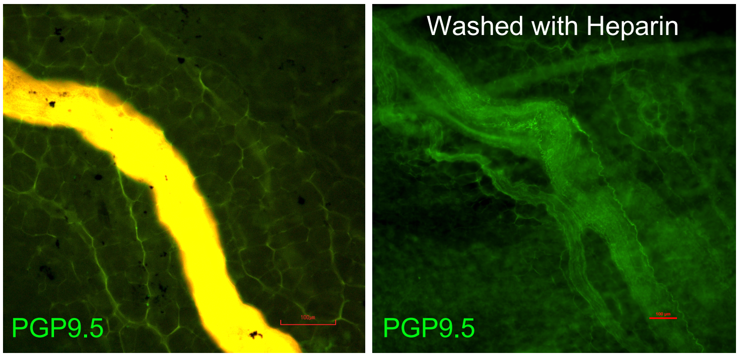


Heparin reduces auto-fluorescence of vasculature

**Supplemental Data**

#### Supplemental Figure S7 – Neuronal reporters for specific nerve subtypes compared to the respective antibodies

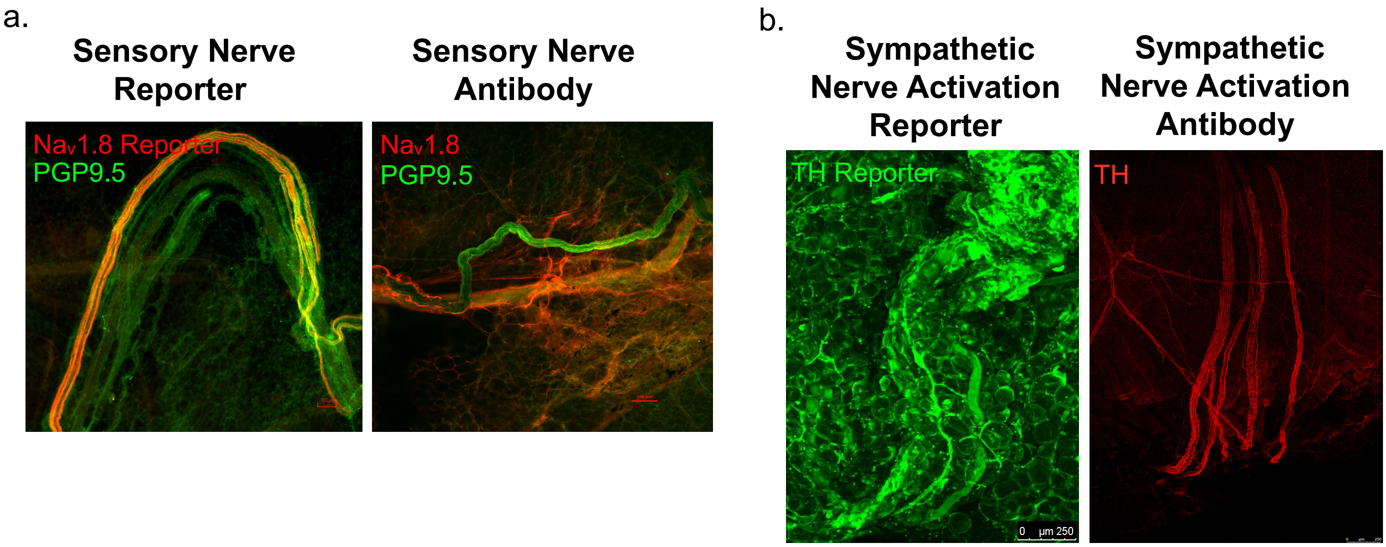

**Supplemental Data**

**Supplemental Figure S8 – AAV mediated BDNF delivery to inguinal scWAT of BTBR MUT mice**

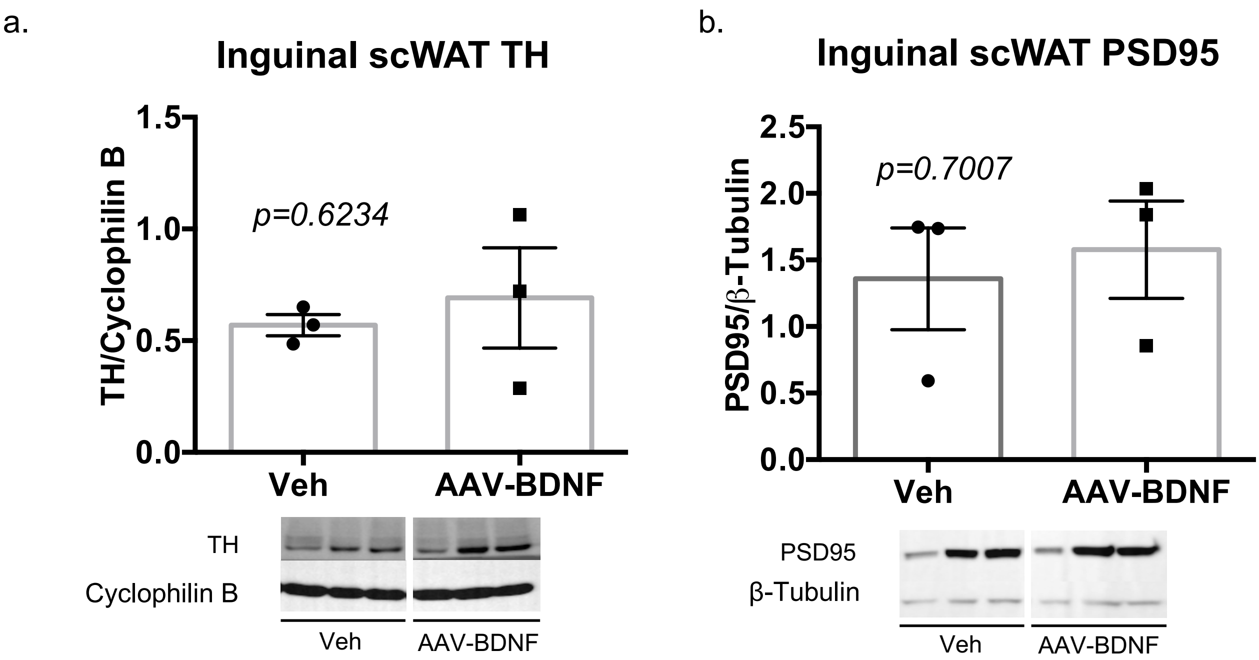

#### Supplemental Data

#### Supplemental Figure S9 – Neurovascular interactions

a. **Inguinal scWAT Luxol Blue Myelin Stain**

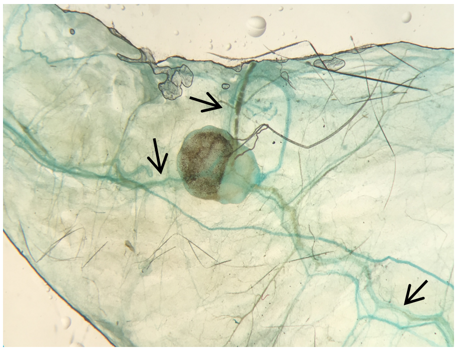

**Digital zoom of 30x mag**

b. **Inguinal scWAT Innervated Vasculature**

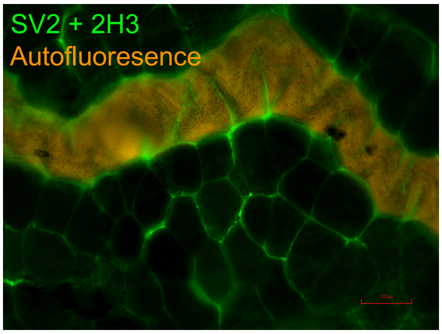


c. **Neurovascular Interactions in Inguinal scWAT**

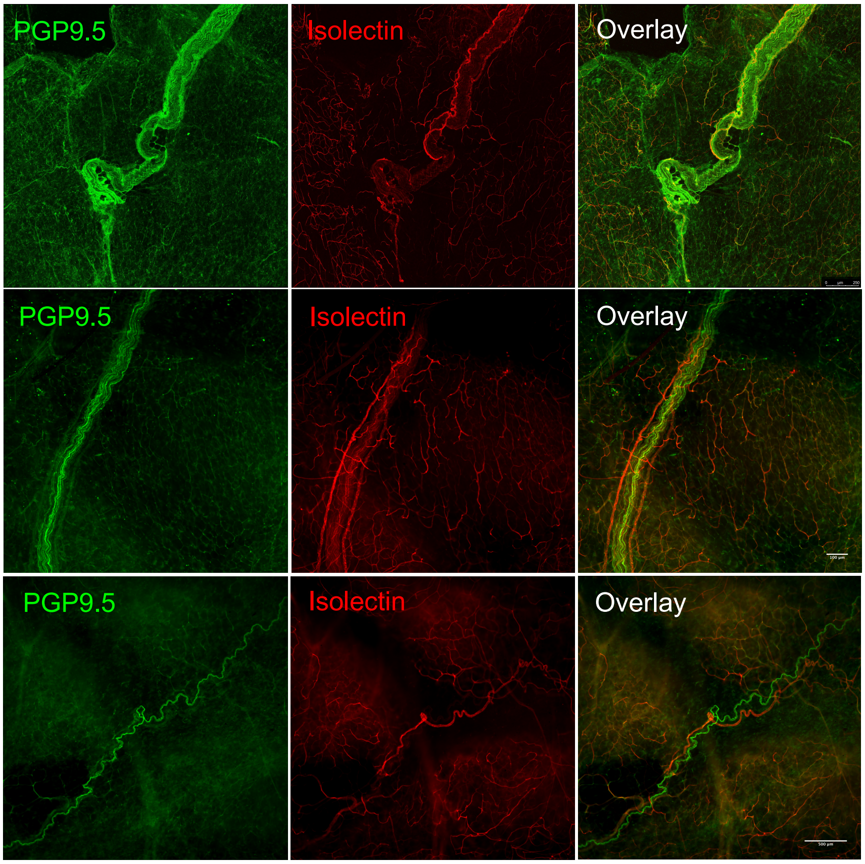

#### Supplemental Data

#### Supplemental Figure S10 – Adipose innervation: Sex and depot comparison

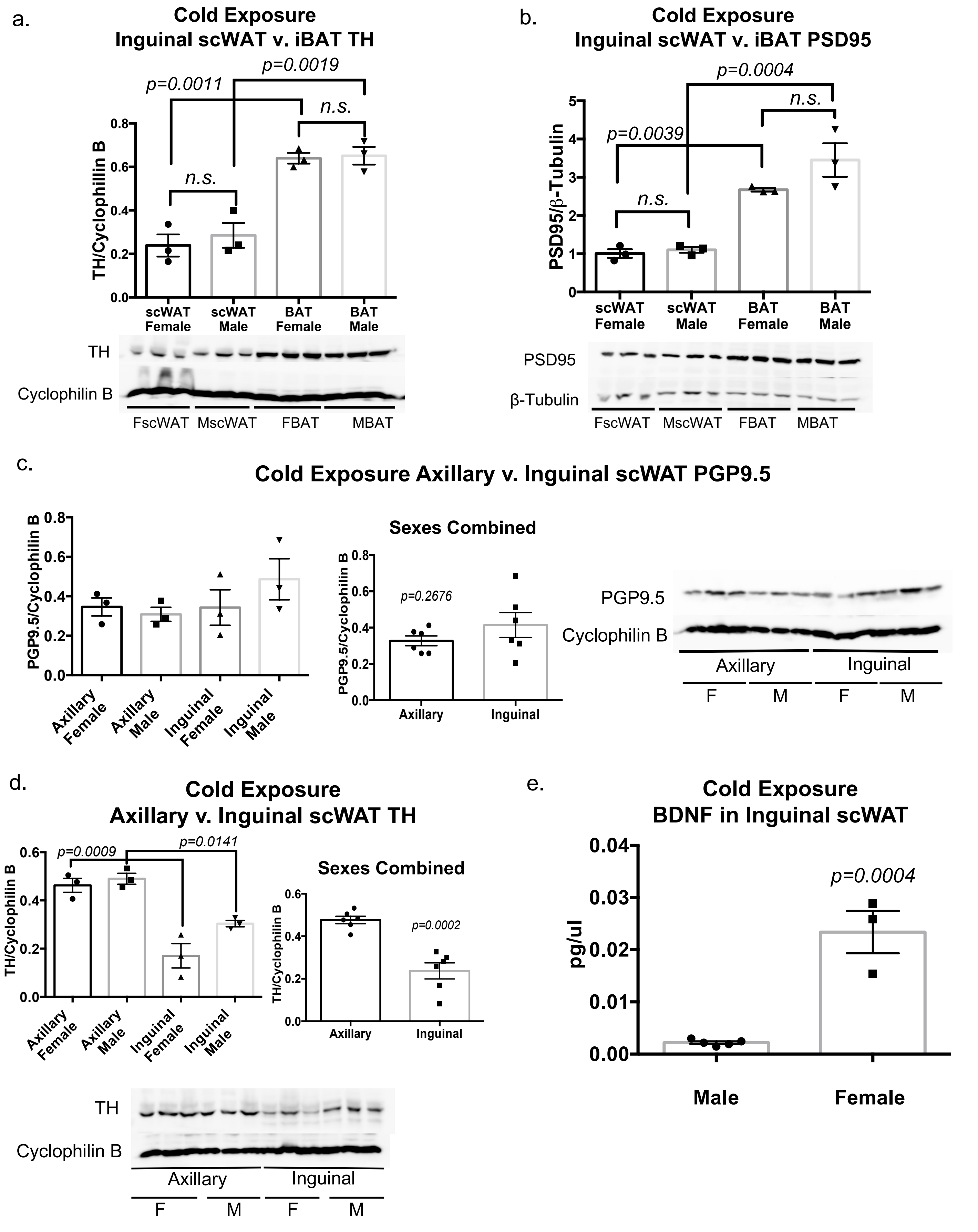
